# Repeated introduction of micropollutants enhances microbial succession despite stable degradation patterns

**DOI:** 10.1101/2021.08.24.457489

**Authors:** Dandan Izabel-Shen, Shuang Li, Tingwei Luo, Jianjun Wang, Yan Li, Qian Sun, Chang-Ping Yu, Anyi Hu

## Abstract

The increasing-volume release of micropollutants into natural surface waters has raised great concern due to their environmental accumulation. Persisting micropollutants can impact multiple generations of organisms, but their microbially-mediated degradation and their influence on community assembly remain understudied. Here, freshwater microbes were treated with several common micropollutants, alone or in combination, and then transferred every 5 days to fresh medium containing the same micropollutants to mimic the repeated exposure of microbes. Metabarcoding of 16S rRNA gene makers was chosen to study the succession of bacterial assemblages following micropollutant exposure. The removal rates of micropollutants were then measured to assess degradation capacity of the associated communities. The degradation of micropollutants did not accelerate over time but altered the microbial community composition. Community assembly was dominated by stochastic processes during early exposure, via random community changes and emergence of seedbanks, and deterministic processes later in the exposure, via advanced community succession. Early exposure stages were characterized by the presence of sensitive microorganisms such as Actinobacteria and Planctomycetes, which were then replaced by more tolerant bacteria such as Bacteroidetes and Gammaproteobacteria. Our findings have important implication for ecological feedback between microbe-micropollutants under anthropogenic climate change scenarios.

## Introduction

The rapid exploitation and widespread use of emerging organic chemicals have resulted in environmental damage and human health issues during the last several decades [1–3]. These organic chemicals include pharmaceuticals and personal care products, hormones, and industrial chemicals, and are typically classified as micropollutants, because of their occurrence at a low to very low concentration of μg and ng per litter in the receiving waters [4, 5]. The majority of micropollutants originate from direct and indirect inputs of organic contaminants that are released via the effluents of wastewater treatments plants [1, 6–8]. The micropollutants gradually accumulate at sites of their release and reach concentrations high enough to exert disruptive effects on aquatic organisms such as fish, invertebrates, and microbes [9–13]. Nonetheless, little is known about the mechanisms by which micropollutants impact the turnover of aquatic microbial communities [7, 14, 15], especially those of freshwater bacteria [16, 17], and the role of microbes on trace micropollutant degradation.

How microbial assemblages respond to micropollutants can be classified as: subsidy responses and stress responses. The subsidy responses is mainly via biodegradation, in which the microbes are able to utilize the micropollutant compounds as carbon sources, while the stress responses is relevant to the growth and survival of microorganisms inhibited [18, 19]. For instance, bisphenol A (BPA) and its substituted derivatives (e.g., bisphenol S, BPS) [20] are typical subsidy responses, as their relatively simple chemical structure and low toxicity allow both to be utilized as carbon sources by microbial isolates [19, 21]. Stress responses are elicited by pharmaceutical and personal care micropollutants such as triclosan (TCS) and its substituted derivatives (e.g., triclocarban, TCC) [22], because both inhibit the growth of several microbial taxa, consistent with their resemblance to antimicrobials [18, 23]. Both BPA and TCS in the field strongly influence microbial diversity, especially that of the central species and module communities comprising the microbial co-occurrence network [24].

Microbial responses to chemical pollutants can be modified by the exposure legacy of the community. For example, microbial communities with a history of containment exposure may differ from contaminant-naïve communities in the rate and extent of chemical degradation [25–27]. They may also differ in their plasticity [14] upon subsequent exposure to similar contaminants. However, the ecological drivers of community succession following repeated micropollutant exposure are not well-understood. Studies of community dynamics over multiple generations and a time period relevant for community stability could provide insights into the ecological traits that determine taxon sensitivity and tolerance, and therefore the ability of microbial community members to degrade micropollutants.

Press disturbances in natural environments can impact multiple generations of organisms [28] by altering the affected ecosystems beyond the possibility of recovery or by causing a shift to a community with a new stable state [29]. Following a stress event such as repeated introduction of micropollutants in recipient environments, communities rely on reassembly from the local seedbanks such that succession eventually results in a new steady state. During the early stages of community dynamics, no-longer-dormant local species undergo a secondary succession characterized by new abiotic and biotic interactions. The emergence of resuscitation from dormancy is considered as stochastic processes that lead to divergent communities [30–32]. When abiotic and biotic conditions allow a stabilization of the initial communities, community formation at a later phase of a press disturbance is expected to converge, through the succession of microbial species [33]. In addition, the plasticity of microbial taxa in response to multiple micropollutant exposure has been shown to increase over multiple generations [14], with later-assembled communities shifting to a new stable state that differs from the one before the exposure.

Here, a press disturbance was simulated in a microcosm experiment in which a reservoir bacterial community was transferred into fresh medium containing trace concentrations of typical micropollutants that were refreshed every 5 days. Several different micropollutants are typically present simultaneously within aquatic environments, but whether their interactions lead to effects on microbial communities that differ from those of single micropollutants is unclear [12]. Thus, we also experimentally manipulated single and multiple micropollutants to determine their impacts on taxon coexistence, the regularity of community assembly, and micropollutant degradation ability. We hypothesized that because BPA elicits a subsidy response and TCS a stress response, the removal rate of BPA would be higher than that of TCS, with larger differences in the removal rates by microbial communities exposed to these micropollutants individually rather than in combination with other micropollutants. We also hypothesized that micropollutants degrade and alter the microbial community compositions after exposure. Accordingly, the microbial communities assembled at the initial and late phases of the micropollutant exposure would differ substantially and are primarily driven by either stochastic or deterministic assembly process.

## Materials and Methods Water collection

We selected the bacterial species pools from Shidou Reservoir, which is one of largest sources of drinking water in Xiamen, China [34] and is therefore under governmental protection from pollution originating from household drainages and a nearby dam (Fig. S1). Unlike other contaminated aquatic environments [18, 35, 36], the concentrations of BPA, BPS, TCS, and TCC in Shidou Reservoir (China) are very low (< 1 ng/L), such that the reservoir’s waters provide an ideal system to investigate the influence of persistent micropollutants on an aquatic microbiome. Surface water (100 L) from Shidou Reservoir (24°41’39”N, 118°0’37”E) was collected at 0.5 m depth on June 1, 2017 (Fig. S1). Temperature, pH, conductivity, dissolved oxygen (DO), and dissolved oxygen saturation, measured *in situ* using a HACH HQ40d multi-parameter meter (HACH, Loveland, CO, USA), were 29.0°C, 8.90, 54.4 μs/cm, 8.84 mg/L and 116.7%, respectively. The concentrations of BPA, BPS, TCS, and TCC in the reservoir water were measured by liquid chromatography-tandem mass spectrometry (LC-MS/MS) (ABI 3200QTRAP, Framingham, MA, USA) with solid-phase extraction and all were < 1 ng/L at the time of sampling (details of the measurements are provided below). The collected water was transported from the field to the laboratory within 2 h after sampling.

### Experimental setup

BPA, BPS, TCS, and TCC were dissolved in acetone at concentrations of 800, 800, 400, and 400 μg/L, respectively, and stored as stocks in glass bottles (HPLC grade). The micropollutants were previously shown to be chemically stable when dissolved in an organic solvent [36, 37]. From the 100 L of collected water, 15 L were filtered through a 1.2-μm polycarbonate filter (Millipore, Ireland), to minimize top-down regulation by planktonic grazers, and then used as the initial inoculum. The remaining unfiltered water was filtered through a 0.22-μm polycarbonate filter (Millipore, Cork, Ireland) to remove microorganisms and then stored in the dark for later use in the preparation of incubation medium. Two treatments consisting of single micropollutants (BPA and TCS) and two treatments consisting of micropollutant mixtures (MI = BPA + TCS) and (MII = BPA + BPS + TCS + TCC) were established.

Time-serial microcosms comprising seven inoculation batches from batch 1 (named hereafter ‘B1’) through batch 7 (‘B7’) were established to investigate the microbial community response to a press disturbance of micropollutants resulting from a single short-term event but maintained for 35 days (Fig. 1). Prior to the experimental initiation, the caps of the bottles containing the micropollutants were loosened for > 12 h to volatilize the acetone in a fume hood, as described in [38, 39]. In microcosm B1, 600 ml of the initial inoculum was added to bottles that already contained the micropollutants and nutrients, such that the final concentration of each micropollutant in the treatments was as follows: BPA: 1 μg/L BPA, TCS: 1 μg/L TCS, MI: 1 μg/L BPA + 1 μg/L TCS, and MII: 1 μg/L BPA + 0.1 μg/L BPS + 1 μg/L TCS + 0.1 μg/L TCC. BPS and TCC are the derivatives of the BPA and TCS, respectively, and the concentrations of those analogues in the environment are one order of magnitude lower than their chief compounds in the seawater [20, 40, 41]. The final concentration of each micropollutant in the microcosms closely represented their ambient environmental conditions as reported in literature [18, 42]. Further, given the low nutrient concentrations [34, 43], we adjusted the concentrations of nitrogen and phosphorus in the filtered water to 128.6 and 8.04 μM to promote microbial growth by the addition of KNO_3_ and KH_2_PO_4_, respectively [44]. After 5 days of incubation, 60 mL of the B1 inoculum was pipetted from each of the replicates and transferred to a sterile bottle containing 540 mL of 0.22-μm-filtered water and freshly added micropollutants. This newly inoculated microcosm served as batch 2 (named hereafter ‘B2’). The micropollutant treatments and controls were amended with 128.6 μM N and 8.04 μM P after each inoculation to maintain similar amounts of these nutrients throughout the experiment. The same inoculation procedure was applied at each 5-day interval to initiate microcosms B3–B7, such that the experimental period was 35 days (Fig. 1). Five days was chosen as the inoculation duration because previous unpublished observation (Fig. S2) indicated that reservoir bacteria exposed to a micropollutant-added medium had shown a high removal efficiency in the early growth stage, and that was expected to lead up to stationary phase on day 5 of the culture. Microbial cells were transferred between pairwise replicates, e.g., replicate 1 in B1 was the inoculum for replicate 1 in B2, with the remainder used in the downstream analyses described below. An additional set of experimental bottles without micropollutant addition served as the controls. To maintain the same level of exposure to acetone in all microcosms, the controls also received the same spiking of solvent acetone and were left for overnight evaporation before each transfer. In total, there were five treatments, each with three replicates, for a total of 15 microcosm replicates per batch. All microcosms, i.e., 105 experimental bottles were incubated in the dark at 28°C with shaking at 100 rpm.

**Figure 1.**
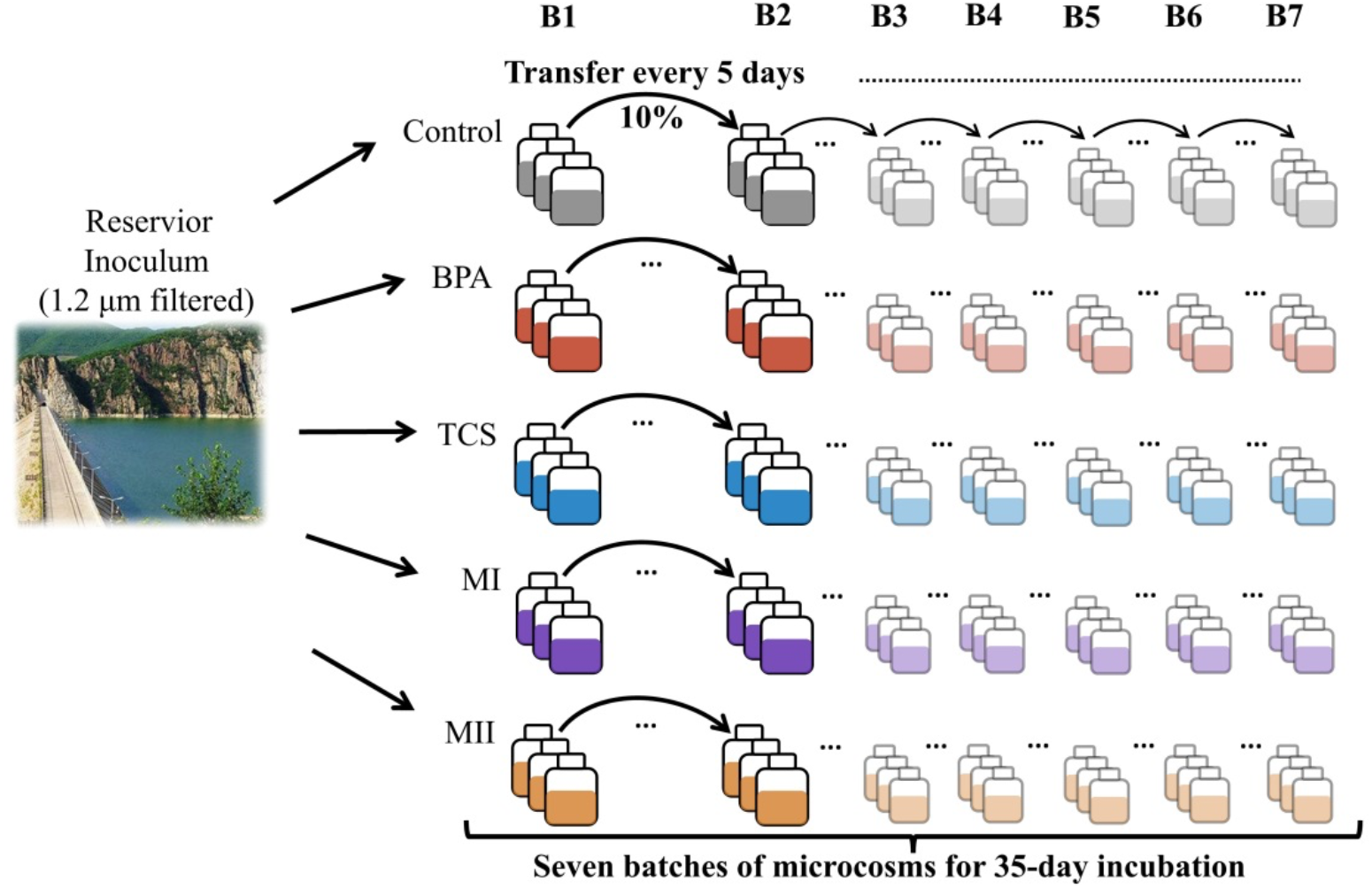
Experimental setup. In microcosm batch 1 (B1), 600 ml of the initial inoculum was added to bottles containing the micropollutants. Starting from the B1, 60 mL (10%) of the inoculum was pipetted from each of the replicates and transferred to fresh medium containing 540 mL of 0.22-μm-filtered water and freshly added micropollutants every 5 days. This procedure was carried out in the seven batches of microcosms from batch 1 (B1) through batch 7 (B7). Samples for bacterial community characterization and micropollutant removal were collected prior to the next inoculation. Treatments are color-coded: ‘Control’ is (without micropollutant addition, grey), ‘BPA’ (microcosms added with bisphenol A, red), ‘TCS’ (microcosms added with triclosan, blue), ‘MI’ (microcosms added with a mixture containing bisphenol A and triclosan, purple), ‘MII’ (microcosms added with a mixture containing bisphenol A and triclosan, bisphenol S and triclocarban, yellow).

Additional experiments were conducted to determine (i) micropollutant removal rates in the absence of microorganisms, to rule out the gradual degradation of the micropollutants in the treatments due to their chemical instability (Fig. S2), and (ii) whether the ability of the microorganisms to transform micropollutants differed depending on their pre-exposure history (Fig. S3). Please see details on these two experiments in the Supplemental methods part 1.

### Micropollutant analysis

For each microcosm batch, ∼20 mL of the inoculum was removed from each bottle at 0, 2 and 5 days for micropollutant analysis. The four micropollutants BPA, BPS, TCS, and TCC were concentrated and purified by a 12-port Visiprep-DL SPE vacuum manifold (Supelco, Bellefonte, PA, USA) with Oasis HLB cartridges (500 mg, 6 mL; Waters, Millford, MA, USA), and then were measured using LC-MS/MS (ABI 3200QTRAP, Framingham, MA, USA), as described in our previous work [37]. Chromatographic separation was performed by using a Kinetex C18 column (100 mm × 4.6 mm, 2.6 μm; Phenomenex, Torrance, CA, USA) from Shimadzu LC system (Shimadzu, Japan).

### DNA extraction and 16S rDNA amplicon high-throughput sequencing

The remaining inoculum (∼470 mL) of each replicate was filtered onto 0.22-μm polycarbonate filters (Millipore, Ireland), which were stored at −80°C until used for DNA extraction. DNA from the 105 samples was extracted using the FastDNA for soil kit (MP Biomedical, Santa Ana, CA, USA) according to the manufacturer’s protocol, except that the beating speed was set to 6.0 m/s for 80 s. The universal primer pair 515F (5’-GTG YCA GCM GCC GCG GTA-3’) and 907R (5’-CCG YCA ATT YMT TTR AGT TT-3’) was used to amplify the V4-V5 region of the bacterial 16S rRNA genes. The PCR products were purified, mixed in equal amounts, and sequenced on an Illumina MiSeq platform with the 2 × 300 bp paired-end protocol at the Majorbio Bio-Pharm Technology Co., Ltd. (Shanghai, China).

### Sequence processing

The raw sequencing data were processed using the LotuS pipeline [45] with the following criteria: i) average sequence quality > 27, ii) sequence length > 170 bp, iii) no ambiguous bases, iv) no mismatches with the barcodes or primers, and v) homopolymer length < 8 bp. Operational taxonomic units (OTUs) were clustered at a 97% similarity using UPARSE [46]. A representative sequence of each OTU was taxonomically quantified using the RDP classifier with the Silva database v132 [47], at a bootstrap cutoff of 80%. All samples were rarified to 36, 000 sequence reads (the smallest library size). The subsampled OTU table contained 1375 OTUs. The α-and β-diversity indices of bacterial communities were calculated from the rarified dataset using QIIME v1.9.1 [48] with the script “core_diversity.py”.

### Inferring the community growth phase from the16S rRNA gene copy number

The 16S rRNA gene copy number among prokaryotes within a defined period indicates the number of individuals within a community and reflects the growth phase of that community [49]. In this study, the 16S rRNA gene copy number of each OTU found in the samples was estimated using the rrnDB database v5.5 [49] and the weighted mean rRNA gene copy number of all OTUs found in each sample was then calculated as described previously [50] (Please see details on the use of rrnDB database and the calculation of weighted mean rRNA gene copy number in the Supplemental methods part 2). The abundance-weighted average gene copy number of a community was plotted against the sampling time for each treatment. The similar overarching pattern of the average rRNA gene copy numbers of bacterial communities in all microcosms (Figs. S4A-F) indicated the similar growth trajectories of those communities over time. A hierarchical cluster analysis with complete clustering criterion and Bray-Curtis dissimilarity metric was used to group the similarity of average gene copy numbers from all micropollutant-treated microcosms belonging to the same batch type, without including the controls. The clustering analyses was done in the R package ‘stats’ package v 3.6.3. Three growth progression phrases across all microcosm batches were identified based on similar average gene copy numbers and the time continuity of the inoculation (Fig. S4G): phase 1 (B1–B2), phase 2 (B3–B4), and phase 3 (B5–B7).

### Relative contribution of ecological processes

According to a quantitative ecological framework [51, 52], community assembly can be summarized by five major ecological processes: i) homogenizing selection for a convergent composition between communities, ii) variable selection for a divergent composition between communities, iii) dispersal limitation for a divergent composition between communities due to limited dispersal events, iv) homogenizing dispersal for a convergent composition between communities due to unlimited dispersal, and v) ecological drift (referred to “undominated processes” in [52]. The phylogenetic turnover among the communities was quantified by the weighted β-mean nearest taxon distance (βMNTD), as described in [53]. The βNTI was calculated as the difference between the observed βMNTD and the mean of the null distribution (999 randomizations). The criteria determining the fraction of each process were as follows: i) homogenizing and variable selection: higher (> 2) and lower (< −2) βNTI, respectively; ii) dispersal limitation and homogenizing dispersal: non-significant |βNTI| (< 2) but RC_bray_ > 0.95 and < −0.95, respectively. iii) ecological drift: non-significant |βNTI| value (< 2) and non-significant |RC_bray_| value (< 0.95) [52]. To infer the relative contributions of the different community assembly processes to community turnover over time, for each pair consisting of a micropollutant treatment and its control the βNTI and RC_bray_ values were calculated separately for each of the three growth phrases described above.

### Ecological grouping of microbial responses to micropollutants

We determined ecological responses of bacterial taxa based on their relative abundances at growth phases (phase 1: B1–B2, phase 2:B3–B4, and phase 3: B5–B7, Fig. S4G) throughout repeated micropollutant exposure, using the ‘DESeq2’ R package v 1.26.0 [54]. Significant values were corrected for multiple tests using the Benjamini-Hochberg correction with an adjusted α value of 0.05. The responses were categorized as: i) sensitive, when relative OTU abundances were significantly higher in phase 1 than in phases 2 and 3; ii) opportunistic, when relative abundances were significantly higher in phase 2 than in phases 1 and 3; iii) tolerant, when relative abundances were significantly higher in phase 3 than in phases 1 and 2. The ecological categories comprising a group of bacterial members whose growth fitness optimal in response to repeated exposure could be documented collectively. In addition, to visualize the phylogenetic relationships of the OTUs assigned to the ecological groups, representative sequences of those OTUs were used to construct a phylogenetic tree using iTOL v5 [55].

### Micropollutant turnover and community analysis

A Wilcoxon’s test was used to assess the removal rate of BPA, TCS, BPS, and TCC in consecutive microcosm batches for the single-micropollutant (BPA and TCA) treatments and the mixed-micropollutant (MI and MII) treatments. Additionally, a Kruskal-Wallis test was used to analyze the difference in the removal rates between the single micropollutant treatments and between any two micropollutants for the mixture treatments. In this case, the removal rate of the micropollutants from each microcosm batch was used as replicates for the analysis.

Differences in the bacterial communities in all microcosms were explored using non-metric multidimensional scaling (NMDS) based on the Bray-Curtis dissimilarity metric and by fitting removal rates of micropollutants to the ordination. A permutational multivariate analysis of variance (PERMANOVA) [56] was conducted to test the effects of micropollutants, inoculation time (as indicated by the microcosm batch), and their interactive effects on the taxonomic composition of the bacterial communities. A pairwise PERMANOVA was performed to test for significant differences in community composition between each treatment pair. The multivariate analyses were performed using the R package ‘vegan’ [57]. Whether communities with the shared taxa exhibit similar dynamics was assessed in a dissimilarity-overlap analysis [58]. Briefly, the relative abundance of each OTU in each community was calculated to identify sets of OTUs within each pair of communities (i.e., within a treatment over time). In the calculation of dissimilarity, only the shared OTUs were considered, and the relative abundance of shared taxa was transformed to add up to 1. The dissimilarity between renormalized vectors X and Y can be calculated using root Jensen–Shannon divergence. This method and the relevant R script were modified from [59] Moreover, the rate of community turnover in relative to the controls over time was quantified with a linear regression model (see the Supplemental methods part 3 for details). A Wilcoxon test was then applied to test for significant differences in community turnover between microcosm B1 (day 5) and the other microcosm batches from later time points (B2– B7), using the ‘ggsignif’ package [60].

## Results

### Transformation of trace micropollutants

There was no evidence that the removal rate of any of the micropollutants differed between any of the consecutive microcosm batches (Fig. 2, Wilcoxon’s test, all *P* > 0.05). In the microcosms treated with single micropollutant, most of BPA added to each microcosm batch was degraded, achieving > 80% removal in the individual batches, whereas the removal rate of TCS was less than 60% (Fig. 2). There was no evidence that the removal of the BPA and TCS differed in the MI microcosms differed (*P* > 0.05) (17.86 ± 15.17% and 25.11 ± 18.88 %, respectively; Fig. S5B). In the MII microcosms, the percentage of removed BPA (65.83 ± 6.54%) was highest (*P* < 0.05), followed by the TCS (13.76 ± 16.76%), BPS (5.93 ± 8.97%), and TCC (11.97 ± 14.08%), but the differences were not significant (Fig. S5B).

**Figure 2.**
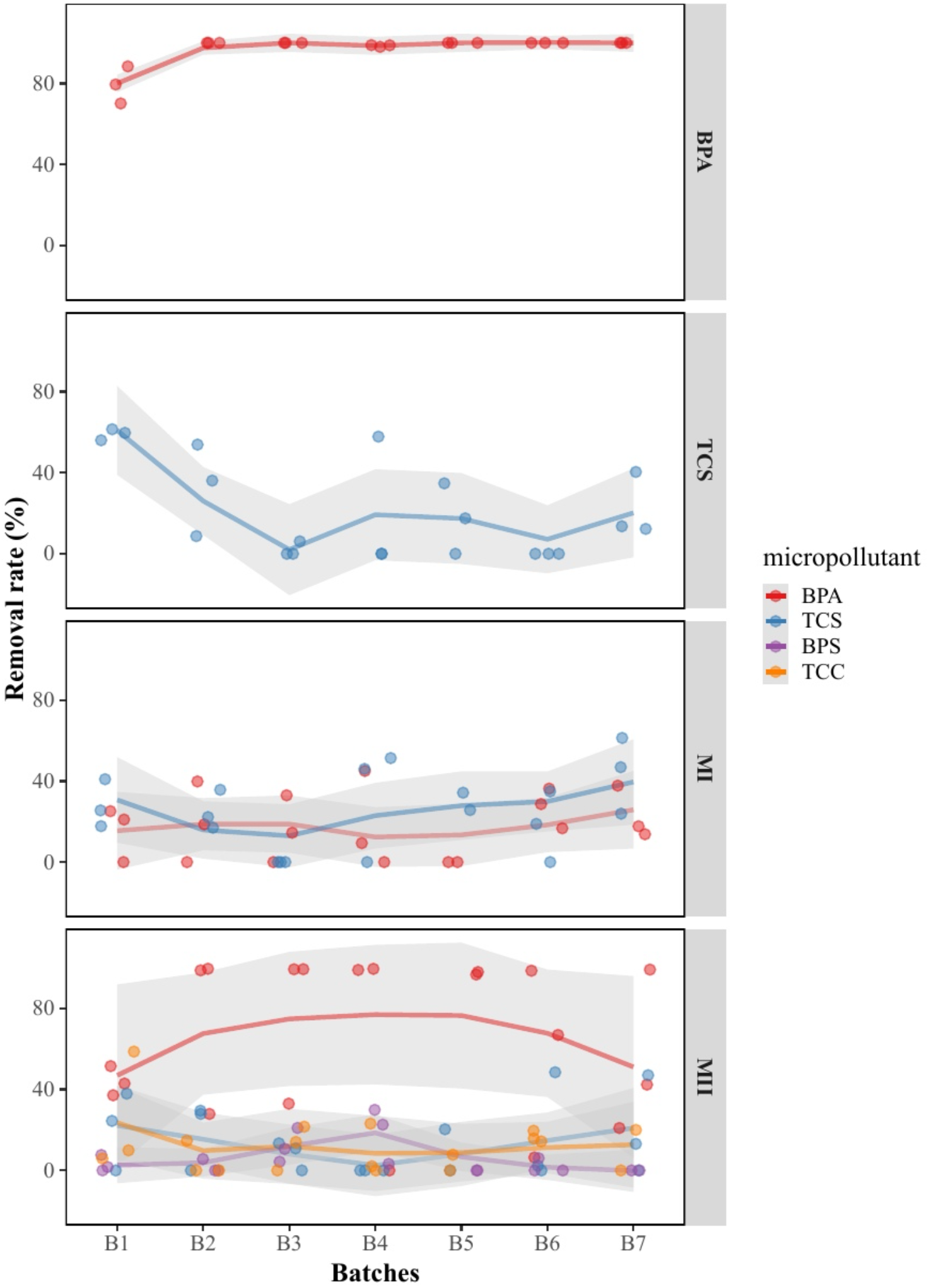
Percent removal of bisphenol A (BPA), triclosan (TCS), bisphenol S (BPS), and triclocarbon (TCC) in all microcosm batches for the single-micropollutant (BPA and TCA) treatments and in the mixed-micropollutant (MI and MII) treatments. Removal was defined as the decrease in the concentration of the chemical relative to the total concentration within each batch of microcosms. ‘B1-B7’ represent the number of microcosm batch that corresponds to every 5-day inoculation.

### Community succession following micropollutant exposure

The bacterial communities in the microcosms became taxonomically distinct both over time and by micropollutant type (Fig. 3). There was very weak evidence that the variation in the community composition among all microcosm batches correlated with differences in the removal rates of BPA (*P* =0.06, Table S1), with high association in its single treatments. However, there was strong evidence that the community variation was correlated with TCS, with high association in TCS single and mixed treatments (Fig. 3 and Table S1, *P* =0.001). The inoculation time (PERMANOVA, R^2^ = 0.149), micropollutant addition (R^2^ = 0.11), and their interaction (R^2^ = 0.055) significantly influenced the community composition (all *P* < 0.001) (Table S2). Irrespective of the inoculation time, the composition of the TCS, MI, and MII communities differed significantly from that of the controls (pairwise PERMANOVA, *P* < 0.01; Table S2) but not from each other. Conversely, the BPA communities significantly differed from the other three treatments but not from the controls.

**Figure 3.**
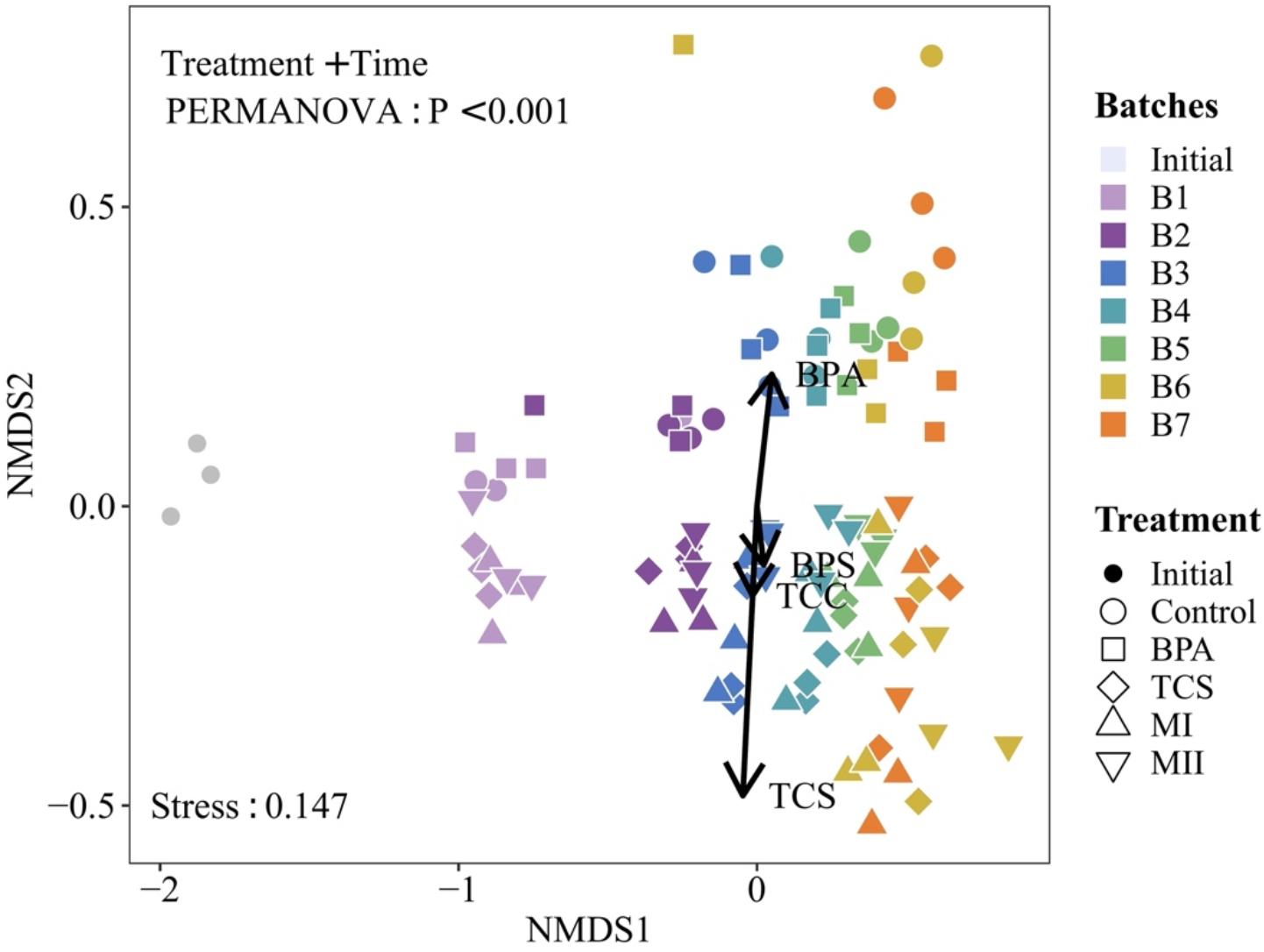
Difference in the bacterial communities in all microcosms batches as determined by non-metric multidimensional scaling (NMDS) ordination. Different treatments are represented by different symbols, and microcosms batches are color-coded. Explanatory environmental variables fitting the ordination are shown with solid arrows. Explanatory values of the environmental variables to differences in the communities are displayed in Supplementary Table S1. Treatment IDs are: ‘Initial’ containing the starting bacterial communities before the experimental implementation; ‘Con’ microcosms containing no micropollutant additions and serve as the controls; ‘BPA’ microcosms containing bisphenol A; ‘TCS’ microcosms containing triclosan; ‘MI’ microcosms containing a mixture of bisphenol A and triclosan; ‘MII’ microcosms containing a mixture of bisphenol A, triclosan, bisphenol-S and triclocarban. ‘B1-B7’ represent the number of microcosm batch that corresponds to every 5-day inoculation. Explanatory values of the environmental variables to differences in the communities are presented in Supplementary Table S1.

We also performed sequence processing with the use of amplicon sequence variants (ASVs) from the DADA2 pipeline [61], and found that overarching patterns of beta diversity were statistically indistinguishable from those observed using OTUs defined at 97% sequence similarity by two separate tests (Fig. S6; Correlation in a symmetric Procrustes M^2^ value = 0.20, *P* < 0.001 on 999 permutations; Mantel correlation = 0.92, *P* < 0.001 on 999 permutations). It has been suggested that broadscale ecological patterns are robust in terms of the use of ASVs vs. OTUs [62]. Therefore, the patterns of microbial community dynamics following repeated exposure to micropollutants can be considered robust regardless of whether 97% OTUs or ASVs are selected.

Overall, the dissimilarity in the micropollutant-treated vs. control communities significantly increased across microcosm batches, as each inoculation was replenished with micropollutants (linear regression, all *P* < 0.001, Fig. S7). Furthermore, the communities of the first inoculation (B1) impacted the response of some communities inoculated at later time points (Fig. 4A). There was no evidence for the impact of the B1 community composition on later communities across B2–B6 in the BPA-treated microcosms and B2–B5 in the MII-treated microcosms, because of their community resemblances (Wilcoxon test, *P* > 0.05; Fig. 4A). In both BPA and MII communities, the correlation between dissimilarity and overlap (i.e., the number of shared OTUs between the community pair) was negative (*P* < 0.05 Fig. 4B). This result was not unexpected since the dispersal of microbial cells from one condition to another similar environment may drive community dynamics towards convergence [63, 64]. However, in the TCS and MI treatments, an impact of the initial communities on later ones occurred only in B3, which is supported by the fact that the B3–B7 communities were more dissimilar to B1 (*P* < 0.05; Fig. 4A). The negative correlation of the dissimilarity and overlap analysis was nonsignificant for the TCS and MI communities such that the high overlap did not result in a low dissimilarity (*P* > 0.05; Fig. 4B). This suggests that there were differences in the dynamics of the shared taxa in those TCS and MI communities over time compared to the communities in the other treatments.

**Figure 4.**
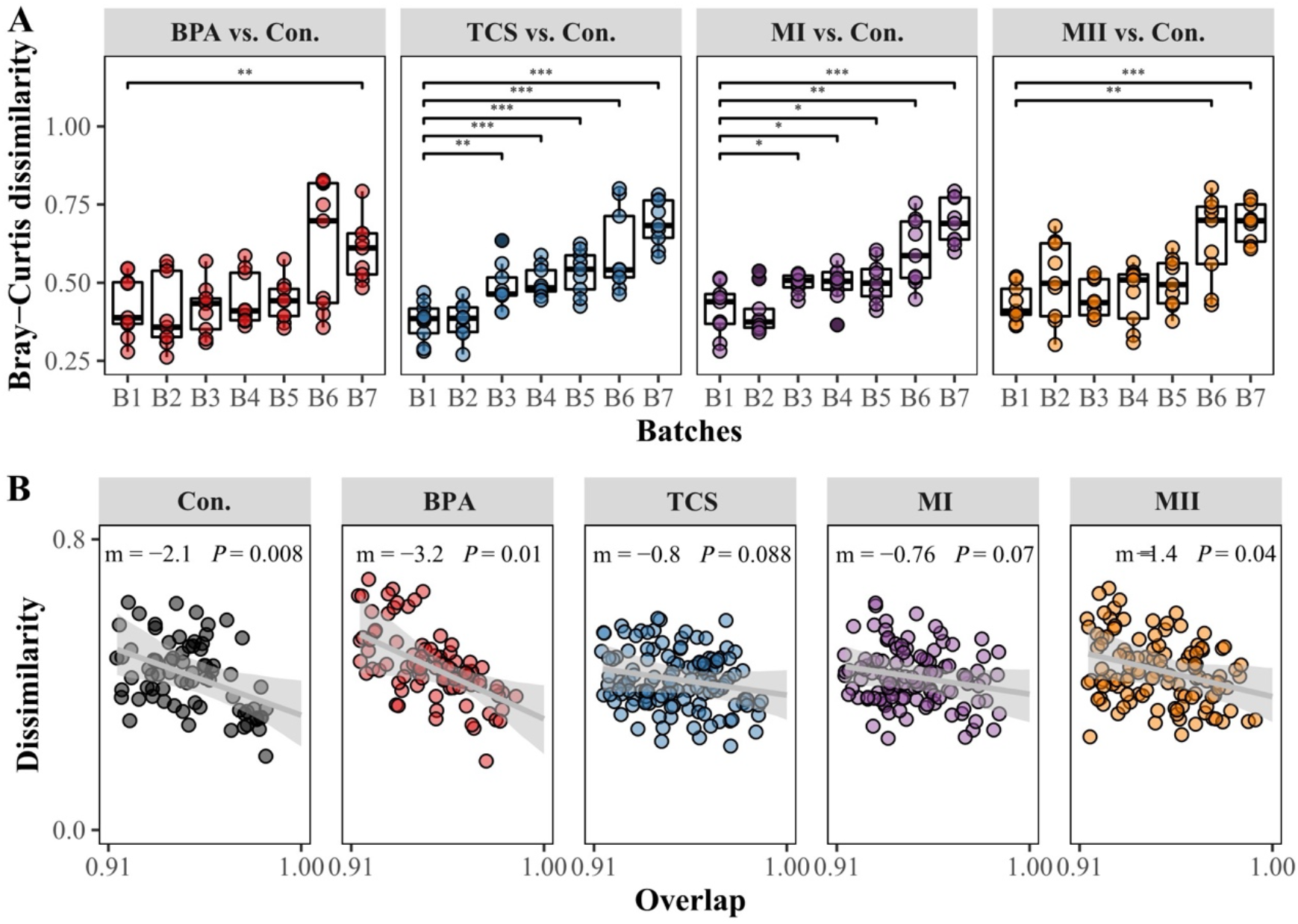
Bacterial community dissimilarity in the micropollutant treatments vs. the controls (i.e., ‘Con.’) over time (A) and the dynamic regularity within treatments (B). Significant differences in the Bray-Curtis dissimilarity for each pairwise comparison of the treatment and control (B1 microcosms vs. later microcosms in (A) were determined in a Wilcoxon test: ***, *P* < 0.001; **, 0.001 < *P* < 0.01; *, 0.01 < *P* < 0.05. The dissimilarity-overlap analysis within each treatment is shown in (B); out of total 210 sample pairs, only the sample pairs that had median overlap above cutoff = 0.913 were shown in the plots. The gray lines indicate the linear regression with the slope (m), and the *P* value (the fraction of bootstrap realizations in which the slope was negative). Gray shaded areas indicate the 95% confidence intervals of the linear regression. Significance level was determined at *P* < 0.05. ‘B1-B7’ represent the number of microcosm batch that corresponds to every 5-day inoculation.

### OTUs with distinct responses to micropollutant exposure

In all treatments, a greater number of sensitive OTUs (41–64), a moderate number of tolerant OTUs (6–15), and fewer opportunistic OTUs (0–3) were identified across phases 1 (B1−B2), 2 (B3−B4), and 3(B5−B7) (Table 1; Fig. 5). The ecological groups replaced one another over time, together accounting for a minimal relative abundance of 0.5 % and a maximum of 60% in the individual samples (Fig. S8). Regardless of the micropollutant type, members of Actinobacteria, Planctomycetes, and Verrucomicrobia primarily consisted of sensitive OTUs (Table 1; Fig. 5). The tolerant OTUs were primarily comprised of members affiliated with Alphaproteobacteria, Gammaproteobacteria and Bacteroidetes. The majority of the tolerant OTUs were unique to one treatment, with only two OTUs (OTU_48 and OTU_78) tolerating micropollutant addition in all treatments (Fig. 5; Table S4). By contrast, many sensitive OTUs (51) were shared between treatments, and at least between two treatments (Table S4). Among the overlapping, 20 OTUs found in all treatments, maximal relative abundance of which was mainly detected in the initial inocula or the controls, and most were members of Actinobacteria (Table S4; Fig. 5). For instance, strictly sensitive taxa Actinobacteria *ACK-M1*, found in both single and mixed micropollutant treatments, exhibited its maximal relative abundance (15.94%) in the initial inocula. Whereas a tolerant taxa Alphaproteobacteria *Novosphingobium*, shared between TCS and MI, had its maximal relative abundance (30.48%) in the BPA treatments at late exposure batches (B6), which has been reported as TCS-degrading bacteria in wastewater treatments and agricultural soil [65, 66].

**Table 1.** Number of OTUs assigned to the ecological categories opportunistic, sensitive, and tolerant in the four micropollutant treatments: ‘BPA’ microcosms containing bisphenol A, ‘TCS’ microcosms containing triclosan, ‘MI’ microcosms containing a mixture of bisphenol A and triclosan, ‘MII’ microcosms containing a mixture of bisphenol A, triclosan, bisphenol S and triclocarban.

| Treatment | Number of OTUs in each ecological category |  |  |
| --- | --- | --- | --- |
|  | Opportunistic | Sensitive | Tolerant |
| BPA | 1 | 64 | 6 |
| TCS | 1 | 47 | 15 |
| MI | 3 | 51 | 12 |
| MII | 0 | 41 | 10 |

**Figure 5.**
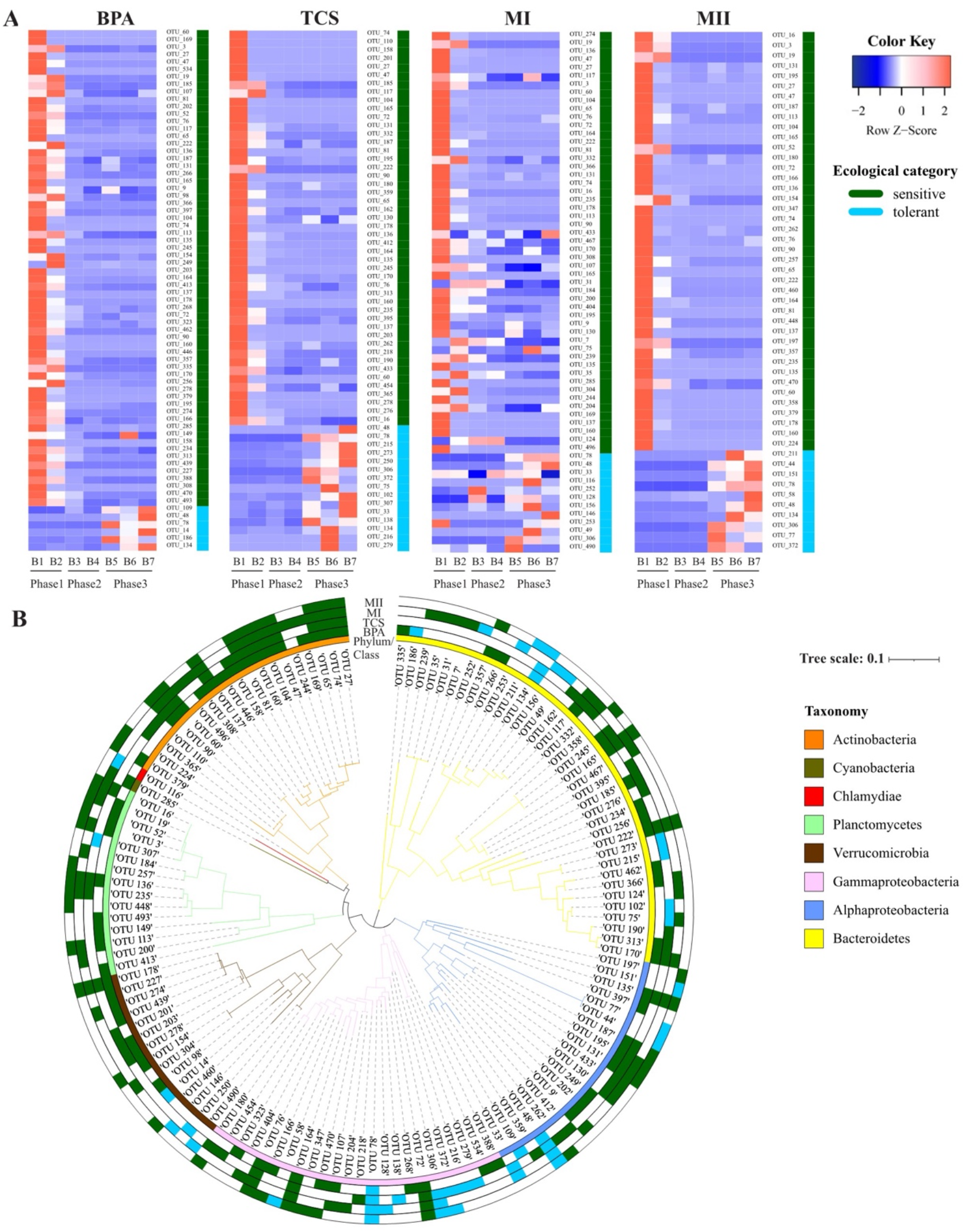
Changes in the relative abundances (rows of heatmap) of the OTUs assigned to the ecological categories tolerant and sensitive across the three growth phases (columns of heatmap) (A) and their phylogenetic distribution (B). In (A), color gradients of heatmap represent the relative abundances of individual by column with warm colors (toward red) indicating high abundance and cold colors (toward blue indicating low abundance within the sample. Column labels are batch IDs and the grouping of growth phases, and row labels the OTU IDs. Tolerant and sensitive OTUs were denoted in green and blue colors, respectively. In (B), the innermost circle indicates the taxonomy (phylum/class) of the OTUs. The outer circles indicate the ecological group of OTUs in each treatment group (BPA, TCS, MI, MII from inside to outside, respectively).

We further checked if the identification of those ecological categories is biased towards the selection of bacterial taxa that were simply enriched in the controls. A large fraction of the bacterial OTUs assigned with ecological strategies (> 67% of the total OTUs surveyed) was unique to the micropollutants-added microcosms, which are not shared in the controls when analyzed with the same filtering criteria as done for ecological grouping (Fig. S9). This suggests that OTU ecological assignment was robust when accounted for bias presented by composition data derived from the controls, although there was little overlap in taxon responses between the controls and the treatments.

### Partition of ecological processes on community variation

Overall, homogenizing selection played a minor role in influencing community variation across phases for each community pair (micropollutant-treated and control), as it accounted for less than 8.33% of the ecological processes (Fig. 6). Instead, variable selection (13.89–61.73%) and ecological drift (30.86–80.56%) dominated the ecological processes across the three phases. These two processes exhibited compensatory contributions, showing increasing variable selection but decreasing ecological drift along the phases (Fig. 6). In the case of dispersal limitation, its relative proportion was greater at phase 1 than the other phases for all treatments, especially BPA and MII.

**Figure 6.**
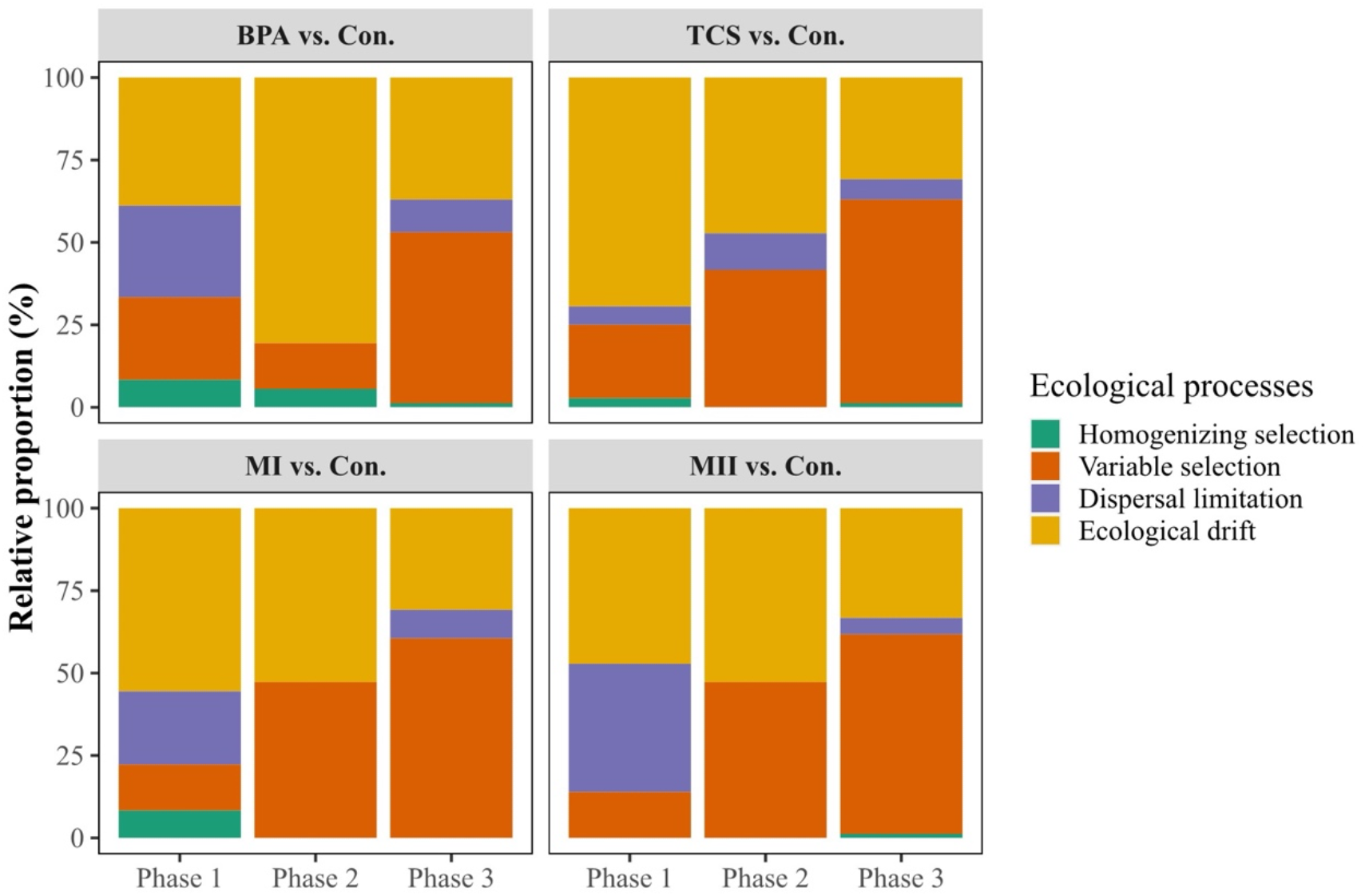
Relative proportion of the ecological processes (homogeneous selection, variable selection dispersal limitation, and ecological drift) contributing to the communities assembled in the micropollutant treatments (BPA, TCS, MI or MII) vs. the controls (i.e., ‘Con.’). The relative contribution was expressed as the portion of pairwise sample assembly that could be attributed to each ecological process. Each of the three phases (corresponding to the early, mid, and late stages of the press disturbance) was classified according to the similarity in the average 16S rRNA gene copy numbers as an indicator of community growth progression. Three growth phrases across microcosm batches were: phase 1 (B1–B2), phase 2 (B3–B4), and phase 3 (B5– B7), as shown in Fig. S4G.

## Discussion

We experimentally investigated the effects of persistent micropollutants on microbial community dynamics to gain insights into community assembly mechanisms. Support was mixed for our first hypothesis that differences in the removal rates of BPA vs. TCS in environments exposed to single vs. mixed micropollutants. Although we observed a certain degree of degradation for each of the four micropollutants, the removal rates of each micropollutant differed depending on its presence alone or with other micropollutants (Fig. 2 and Fig. S5). The BPA degradation was much greater than that of TCS when the microbial communities were exposed to them separately (Fig. S5), suggesting that the former elicits a subsidy response and the latter a stress response on microorganisms [19, 21]. In contrast, these distinctive responses were counterbalanced in the MI treatment, in which the microbial communities were exposed to a mixture of BPA and TCS and there was no evidence found for the differences in their removal rates. The presence of TCS may have inhibited the growth of bacteria that are capable of degrading BPA. As such, the energy required by the microbial communities to counteract the opposing responses to BPA and TCS in the MI treatment likely accounted for the profound inhibition of BPA degradation. In MII, a significantly larger fraction of BPA than of the three other micropollutants was removed. We hypothesize that the presence of TCC reduced the inhibitory effect of TCS on BPA transformation and thus degradation, even if the total concentration of TCC was 10-fold lower than that of TCS in the microcosms. This could be tested by examining the removal efficiency of BPA in the presence of TCC vs. that of both TCC and TCS with varying quantity, but overall this finding highlights that the importance of micropollutant characteristics in understanding the biodegradation potential performed by microorganisms. Moreover, phenolic compounds (both BPA and BPS) act as a growth medium for microorganisms by providing a source of carbon and energy. However, they differ in degradability in the seawater (BPA > BPS)[67], as well as negative impacts on biochemical and microbiological activity of soil (BPA < BPS) [68]. These lines of reasoning explained a greater degradation of BPA than its analogue in MII (Fig. 2). The removal rates of micropollutants measured in our study are regarded as the potential rather than absolute removal efficiency. This is because we cannot rule out a potential role of labile carbon sources present in the 0.22-μm filtered medium in influencing microbial capacity on degrading micropollutants. However, the starting concentration of labile carbon sources present in all media was almost constant as they originated from the same filtered freshwater. Thus, any effects of labile carbon sources on the bacterial communities were most likely consistent in all the microcosms.

Our additional experiments showed that transformation of the micropollutants is negligible in the absence of microorganisms and light (Fig. S2). Microorganisms pre-exposed to low concentrations of BPA and TCS fully degraded higher concentrations of those micropollutants during subsequent exposures and degradation occurred at a faster rate (Fig. S3), but only when BPA and TCS were provided alone and not in combination. Overall, pre-exposure to micropollutants can enhance the degradation capacity of microbes later confronted with those or similar chemicals, which is consistent with previous literature [14, 27].

Given that the ability of bacterial communities to transform the four micropollutants surveyed, the questions remain: whether micropollutants accelerate degradation through changes in bacterial communities, or the micropollutant degraded and then altered the community composition? In our microcosms, each transfer was random with respect to taxon identity, and such that no particular taxa were favoured or disfavoured in one batch before being dispersed to the new batch. The variation between the replicate batches was due to stochasticity in community assembly, resulting in a slight but nonsignificant deviation in degradation patterns of micropollutants over time. We found no evidence that the degradation of any micropollutants differed across microcosm batches. Rather, there was strong evidence on the differences in the bacterial communities between the micropollutant-added microcosms vs. the controls, and that community turnover increased over time following micropollutant exposure (Figs. 4A, S7). These results supported the second hypothesis that the degradation of micropollutants alters the microbial communities. Such community dynamics is mainly linked by the degradation of BPA and TCS in the microcosms. We identified ecological groups based on changes in the relative abundance of taxa following repeated micropollutant exposure, which integrates population growth, death, survival, and reproduction over 35 days. Both sensitive and tolerant taxa identified here may have affected micropollutant degradation either directly or indirectly, such as by providing enzymatic repertoires to other community members that enabled degradation, including interspecies interactions [69, 70]. The larger proportion of sensitive than tolerant taxa in all treatments suggested that the microbial community was not robust to the persistence of micropollutants in its surroundings. Noteworthily, two strictly tolerant genera across treatments were *Sphingomonas* and *Hydrogenophaga* (Table S4), which have been shown to be enriched in the BPA or TCS-contaminated environments [68, 71, 72] and to produce enzymes capable of degrading different pollutants [73].

Our results supported the third hypothesis that communities assembled at the early stage of micropollutant exposure would be compositionally distinct from those that formed at the later stage. Our findings generally corroborated previous studies showing that the persistence of micropollutants even at low concentrations can influence microbial community membership and abundance over time [24, 27, 74]. Despite a substantial community variation between initial and late stages of the exposure, community trajectory in the BPA and MII were driven by a set of shared taxa that interacted similarly with the respective environments and other community members. Conversely, the less pronounced negative dissimilarity-overlap relationships in the case of the TCS and MI treatments indicated that the microbial taxa shared among different inoculation batches within those treatments interacted differently with the same environment over time. Hence, microbial communities assembled in identical habitats can perceive those habitats differently such that their feedback on the surroundings and other, coexisting members will differ as well, as noted elsewhere [59, 75].

We observed a gradual transition from stochastic to deterministic processes towards the later stage of micropollutant exposure. In all treatments ecological drift played a major role during phases 1 and 2 of community growth progression. This can be explained by the random changes in abundances within microbial communities exposed to micropollutants in confined microcosms, where dispersal is limited and locally dormant cells are about to be awakened to colonize empty niches in the environments upon disturbances [33, 76, 77]. In fact, sensitive taxa declined in the relative abundances while tolerant taxa had negligible abundance (Fig. S8) during the growth phases 1-2 (B1-B2 and B3-B4). Hence, tolerant taxa identified in this study were recruited from rare member pool at the early stage of exposure and regrew to a large population size (∼ relative abundance of 10 %) at the later stage of exposure. Moreover, during early exposure, microbial communities might have perceived no differences in the environmental conditions between the controls and the treatments as they were exposed under environmental levels of micropollutants. However, the increasing contribution of variable selection to community assembly over time suggested that the high phylogenetic turnover of the communities was driven by heterogeneous abiotic and biotic conditions [52, 78]. The increasing effects of micropollutants on the interactions among community members, such that the drivers of community assembly no longer resembled those of the control and resulted in a high rate of community turnover. Our findings suggest that the increasing differences in the abiotic and biotic conditions imposed by the repeated introduction of micropollutants across the microcosms led to higher species selection over time and thus enhance microbial succession.

We also considered the potential role of priority effects on community assembly, independent of the ecological process model used in our study. Priority effects emerge when early colonizers exhaust or modify available resources, thus influencing the establishment success of later colonizers. A strong priority effect in the case of TCS and MI was evidenced by the significant difference in the composition of the initially inoculated vs. any of the later inoculated communities (Fig. 4A). Once they were re-inoculated, taxa that had thrived in the B1 batch modified the environments of later batches, such that some taxa (presumably activated from a seed bank, given that no source of new taxa was externally introduced into the systems) were less or more favorable in later batches. However, for the BPA and MII treatments, the effects occurred at later stage of the press disturbance, suggesting the increasing strength of priority effects with increasing establishment time, as noted elsewhere [79].

Although our microcosms represent an ideal system which is removed from *in situ* conditions, our experimental findings are valuable in at least three aspects, that is : i) the prediction of microbial responses to persistent micropollutants, ii) the harnessing of ecological principles and microbiomes in pollution management [80–82], and (iii) the development of a conceptual model to be used to predict when and how stochastic and deterministic processes contribute to community assembly after the exposure of persistent micropollutants. Our results suggest that strong ecological drift drives random community changes at the early stage of the exposure (Fig. 7). Prolonged ecological drift, variable selection resulting from the effects of micropollutants, and the abiotic and biotic conditions experienced differently by different microbes then determine community succession at the mid-stage of the exposure. Additionally, the high environmental heterogeneity together with the low densities of the communities permitted the emergence of priority effects, in which the early establishment of taxa in micropollutant-containing environments allowed a faster community succession. This was supported by the decrease in the average gene copy number at the intermediate phase (Fig. S4), when community assembly was driven by various ecological drivers [83]. Finally, strong variable selection accompanied by increased priority effects likely set the communities on distinct trajectories, with multiple equilibria at the late stage of the exposure. In fact, more advanced community succession and multiple equilibria characterized the end of the press disturbance, and both played a larger role in the TCS and MII (and to a lesser extent in the BPA) communities than in the MI community (Fig. 7), suggesting convergent recovery of those freshwater bacterial communities to repeated micropollutant exposure.

**Figure 7.**
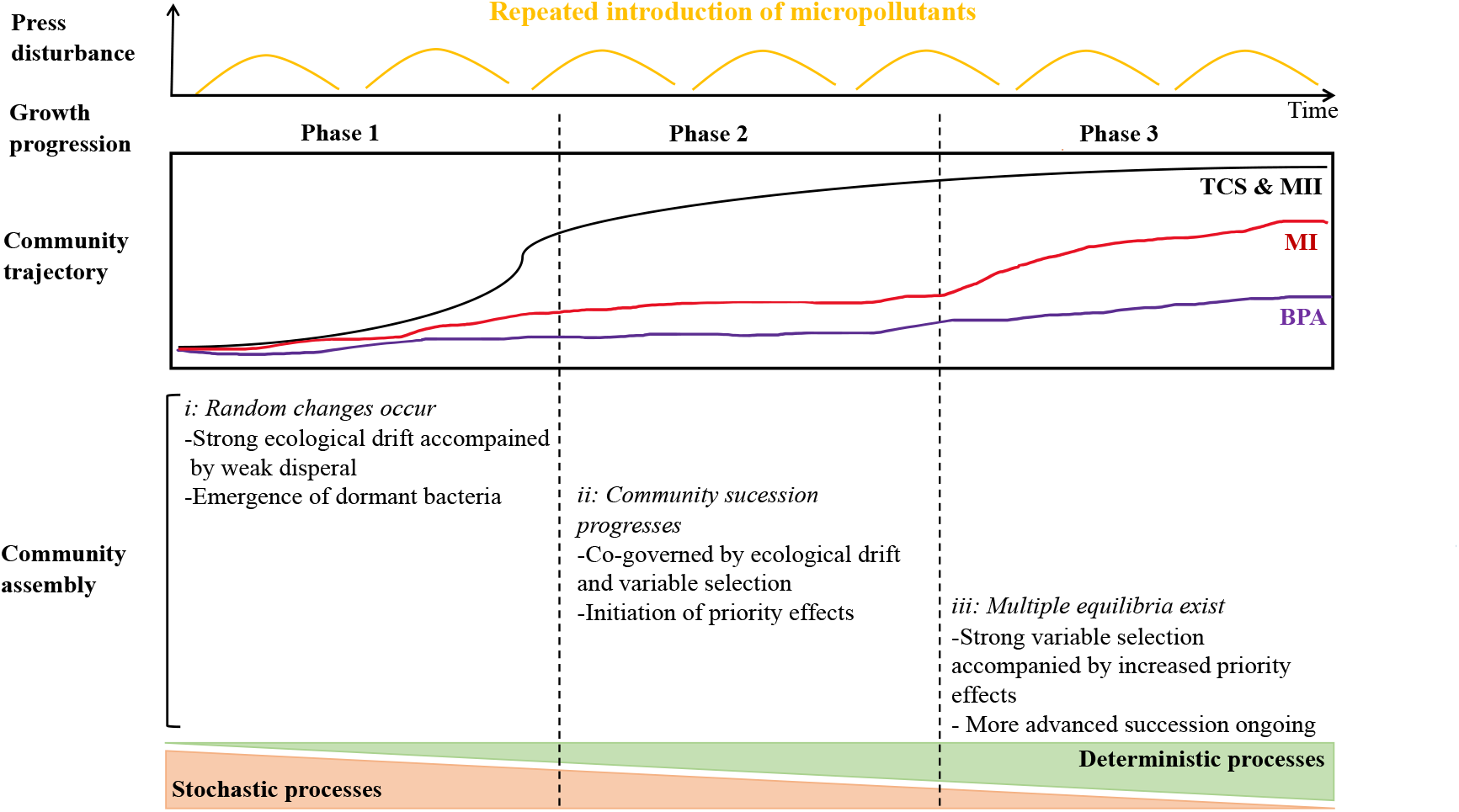
Conceptual model of microbial community assembly processes following a press disturbance. Considering the time continuity of the batch inoculation, the three phases of community growth progression (1, 2, and 3) also correspond to the early, mid, and late stages of the 35-day disturbance. This conceptual model illustrates when and how stochastic/deterministic processes contribute to the dynamics of microbial communities repeatedly exposed to micropollutants. Changes in community trajectory over time in the TCS or the MII treatment are indicated by the black line, MI treatment by the red line, and the BPA treatment by the purple line. Overall, stochastic and deterministic processes act in concert during community dynamics: stochastic assembly dominates at the early stage of a press disturbance/community development, and deterministic assembly at the later stage. Major community assembly mechanisms are delineated for each phase. Community succession accelerates at phase 2 such that multiple equilibria occur at phase 3. Community succession and multiple equilibria are not exclusive, but community turnover is high towards the end of the disturbance.

In summary, we evidenced that micropollutants degrade and drive microbial succession following repeated exposure. Mobile genetic elements (MGEs) function as genetic rearrangement in microorganisms, thus allowing microbial communities to tolerate or adapt to environmental disturbance, such as the presence of organic pollutants [84, 85]. Future work that considers quantification of MGEs using qPCR at successive time points when a microbial community is exposed to micropollutants, would help assess the contribution of MGEs expression to changes in microbial composition and degradation capacity. From the perspectives of microbiome engineering, this study demonstrates the importance of the microbiome’s preexposure history in the design of long-term experiments that include artificial selection to direct the evolution [81, 86] of a target microbiome highly optimized for pollutant removal in an aquatic environment.

## Data and code availability

The sequencing data were deposited in the NCBI SRA database with the Bioproject number PRJNA497759. Our R scripts for statistics, data visualization and computing notes are available on the Github repository (https://github.com/AnyiHu/Impacts-of-micropollutants-on-freshwater-microbiome) and citable using https://doi.org/10.5281/zenodo.5242686.

## Acknowledgements

We thank Dr. Linwei Wu for her assistance in calculating the copy number of 16S rRNA genes, and Dr. Hongjie Wang for his assistance in the molecular experiments. This work was supported by the National Natural Science Foundation of China (31870475 and U1805244), the International Partnership Program of Chinese Academy of Sciences (132C35KYSB20200012), and the Natural Science Foundation of Fujian, China (2019I0030).

## Authors’ contributions

AH and CPY conceived and designed the study. LS and TL performed the experiments and measurement analyses. YL and QS provided the assistance for the LC-MS/MS analysis. DIS, LS, JW and AH performed the data analyses and wrote the manuscript. All authors discussed the results and commented on the manuscript.

## Competing interests

The authors declare no competing interests.

## Notes

### Competing Interest Statement

The authors have declared no competing interest.

### Summary of Updates

This version of the manuscript has been revised to update the methods and discussion section with a few revised figures.

