## Supplemental Information for "Repeated introduction of micropollutants enhances microbial succession despite stable degradation patterns"

**Running title:** Ecological feedback between microbe-micropollutants

Dandan Izabel-Shen[1#], Shuang Li[2,3,4#], Tingwei Luo[5], Jianjun Wang[6],

Yan Li[2,3], Qian Sun[2,3], Chang-Ping Yu[2,7], Anyi Hu[2,3*]

1. Department of Ecology, Environment and Plant Sciences, Stockholm University, 106 91, Stockholm, Sweden
2. CAS Key Laboratory of Urban Pollutant Conversion, Institute of Urban Environment, Chinese Academy of Sciences, Xiamen 361021, China
3. University of Chinese Academy of Sciences, Beijing 100049, China
4. Department of Environmental Microbiology, UFZ, Helmholtz Centre for Environmental Research, Leipzig, Germany
5. Institute of Marine Microbes and Ecospheres, State Key Laboratory of Marine Environmental Science, Xiamen University, Xiamen, China
6. State Key Laboratory of Lake Science and Environment, Nanjing Institute of Geography and Limnology, Chinese Academy of Sciences, Nanjing, 210008, China
7. Graduate Institute of Environmental Engineering, National Taiwan University, Taipei 106, Taiwan

#The authors contributed equally to this work.

*Correspondence to:

Dr. Anyi Hu,

**Contents:**

| | |
|---|---|
| Supplemental text | |
| Part1) Additional experiments to test micropollutant degradation | Page 3 |
| Part2) Calculation of the weighted mean rRNA gene copy number | Page 4 |
| Part3) Analysis of line regression model | Page 5 |
| Supplemental Tables | |
| Table S1 | Page 6 |
| Table S2 | Page 7 |
| Table S3 | Pages 8-14 |
| Table S4 | Pages 15-18 |
| Supplemental Figures | |
| Figure S1 | Page 19 |
| Figure S2 | Page 20 |
| Figure S3 | Page 21 |
| Figure S4 | Page 22 |
| Figure S5 | Page 23 |
| Figure S6 | Page 24 |
| Figure S7 | Page 25 |
| Figure S8 | Page 26 |
| Figure S9 | Page 27 |

### Supplemental methods

#### Part 1) Additional experiments to test micropollutant degradation

An additional experiment was conducted to test the removal rates of micropollutants in the absence of microorganisms. Reservoir water was filtered through 0.22-μm filters and then amended with a mixture consisting of 1 μg BPA/L, 0.1 μg BPS/L, 1 μg TCS/L, and 0.1 μg TCC/L as a blank control. A second additional experiment was conducted to assess whether the ability of prokaryotic assemblages to degrade BPA and TCS was enhanced by the evolving community succession following micropollutant addition. Thus, 40 ml of the initial inoculum and 40 ml of the bacterial assemblage at the end of the experiment (after incubation of the B7 microcosms) were each amended with 160 ml of BPA, 160 ml of TCS, or a mixture thereof, with each micropollutant present at a final concentration of 1 mg/L. The removal rates of BPA and TCS in these treatments after 7 days were measured and the rates in the initial inocula vs. those in the B7 microcosms were compared. The incubation conditions were the same for each treatment and the experiment was conducted in triplicate.

The first experiment showed that the removal proportion of the four micropollutants in the prokaryote-free controls was stable during the 7-day incubation (Supplemental Fig. S2), and therefore the chemical stability and negligible transformation of the micropollutants in the absence of microorganisms. The second experiment showed that the concentration of BPA declined later in the initial community without pre-exposure than in the community with a 35-day history of exposure (Supplemental Fig. S3). In the case of TCS, the concentration did not change over time in the initial community, while in the pre-exposed community it decreased sharply at day 4 of the incubation. These results demonstrated that pre-exposure to micropollutants impacted the ability of the bacterial community to transform micropollutants during a later exposure and that this ability differed between communities exposed to BPA and TCS.

**Part 2) Calculation of the weighted mean rRNA gene copy number**

The rrnDB database catalogues the 16S rRNA gene copy number of an organism with its NCBI or RDP taxonomy [1]. The 16S rRNA gene copy number of an OTU is estimated to be the number of its assigned genus. If the 16S rRNA gene copy number of the corresponding genus is not provided in the rrnDB database, then number of the higher phylogenetic level (i.e., family/order/class/phylum) is used. The abundance-weighed average gene copy number $N_{rrn}$ of each community is calculated using the relative abundance $P$ of each OTU in the community, the taxonomic assignment of each OTU, and the estimated 16S rRNA gene copy number $n$ of each OTU based on rrnDB database v5.5 [2]:

$$N_{rrn} = \sum_{i=1}^{x} (n_i \times P_i)$$

where $n_i$ is the estimated 16S rRNA gene copy number of OTU $i$, $P_i$ is the relative abundance of OTU $i$, and $x$ is the total number of OTU in the microbial community.

#### Part 3) Analysis of line regression model

A linear regression model was used to evaluate the rate of community turnover over time:

$$D = vT$$

where $D$ is the Bray-Curtis dissimilarity between the micropollutant treated vs. untreated communities, $T$ is the incubation time, and the slope $v$ is the turnover rate. The time-decay relationship (TDR) of the PTC of each treatment group and control was evaluated using a log-transformed power law model [3,4]:

$$\log (S) = c + w \log (T)$$

where $S$ is the pairwise Bray-Curtis similarity in community composition across time interval $T$, and the slope $w$ is the temporal turnover rate of the community within each treatment group or control. The significance of the values of slopes $v$ and $w$ was evaluated by 1,000 permutations, using the R package lmPerm [5]. Significant inter-treatment differences in those values were tested by 1,000 bootstraps, followed by a pairwise t-test with Bonferroni correction [6,7].

**Tables**

**Table S1.** Explanatory values of the environmental variables to differences in the communities.

Explanatory values

| | *NMDS1* | *NMDS2* | $R^2$ | *P value* |
|---|---|---|---|---|
| BPA | 0.22 | 0.98 | 0.05 | 0.06* |
| BPS | 0.22 | -0.97 | 0.02 | 0.59 |
| TCC | -0.1 | -0.99 | 0.02 | 0.29 |
| TCS | 0.1 | 1 | 0.24 | 0.001** |

Significance codes: '**' 0.01, '*' 0.1.

Number of permutations: 999

**Table S2.** The results of Permutational Multivariate Analysis of Variance (PERMANOVA) testing the quantitative effects of treatment (i.e., the controls and the type of micropollutant), incubation time (corresponding to microcosm batches), and their interactions on the bacterial community structures across all samples and between treatment pair. $R^2$ values represent the proportion of variance explaining community variation. Bold $R^2$ and $P$ values represent significance of the variance at $P < 0.05$. Treatment IDs are: 'Con' microcosms containing no micropollutant additions and serve as the controls, 'BPA' microcosms containing bisphenol A, 'TCS' microcosms containing triclosan, 'MI' microcosms containing a mixture of bisphenol A and triclosan, 'MII' microcosms containing a mixture of bisphenol A, triclosan, bisphenol-S and triclocarban.

| PERMANOVA | | | pairwise PERMANOVA | | |
|---|---|---|---|---|---|
| Factor | $R^2$ | *P*-value | Pairs | $R^2$ | *P*-value |
| Incubation time | **0.149** | **0.001** | Con. vs BPA | 0.042 | 0.6 |
| Treatment | **0.11** | **0.001** | Con. vs TCS | **0.108** | **0.01** |
| Incubation time *Treatment | **0.055** | **0.001** | Con. vs MI | **0.1** | **0.01** |
| | | | Con. vs MII | **0.095** | **0.01** |
| | | | BPA vs TCS | **0.077** | **0.02** |
| | | | BPA vs MI | 0.067 | 0.06 |
| | | | BPA vs MII | **0.072** | **0.01** |
| | | | TCS vs MI | 0.038 | 1 |
| | | | TCS vs MII | 0.068 | 0.12 |
| | | | MI vs MII | 0.042 | 0.79 |

**Table S3** The ecological strategies of microbial taxa that responded to repeated introduction of micropollutants exhibited distinct response trajectories. Here, the list of OTUs assigned to ecological groups: "opportunistic", "sensitive" and "tolerant" taxa in each micropollutant treatment.

| Ecological category | Treatment | OTUID | Taxonomy (Phylum/class) | log2(P1/P2) | *P.adj* | log2(P2/P3) | *P.adj* |
|---|---|---|---|---|---|---|---|
| opportunistic | BPA | OTU_129 | Bacteroidetes | -8.06 | 0.000 | 4.58 | 0.026 |
| opportunistic | TCS | OTU_376 | Alphaproteobacteria | -4.67 | 0.001 | 3.41 | 0.008 |
| opportunistic | MI | OTU_53 | Planctomycetes | -5.33 | 0.000 | 3.70 | 0.013 |
| opportunistic | MI | OTU_167 | Proteobacteria | -8.91 | 0.003 | 6.64 | 0.027 |
| opportunistic | MI | OTU_126 | Deltaproteobacteria | -5.09 | 0.019 | 4.44 | 0.044 |
| sensitive | BPA | OTU_60 | Actinobacteria | 26.79 | 0.000 | 11.67 | 0.000 |
| sensitive | BPA | OTU_169 | Actinobacteria | 30.00 | 0.000 | 24.26 | 0.000 |
| sensitive | BPA | OTU_3 | Planctomycetes | 8.65 | 0.000 | 4.47 | 0.001 |
| sensitive | BPA | OTU_27 | Actinobacteria | 12.65 | 0.000 | 12.23 | 0.000 |
| sensitive | BPA | OTU_47 | Actinobacteria | 11.77 | 0.000 | 11.68 | 0.000 |
| sensitive | BPA | OTU_534 | Gammaproteobacteria | 22.90 | 0.000 | 23.05 | 0.000 |
| sensitive | BPA | OTU_19 | Planctomycetes | 10.49 | 0.000 | 5.72 | 0.003 |
| sensitive | BPA | OTU_185 | Bacteroidetes | 5.64 | 0.000 | 5.26 | 0.000 |
| sensitive | BPA | OTU_107 | Gammaproteobacteria | 2.71 | 0.000 | 2.29 | 0.000 |
| sensitive | BPA | OTU_81 | Actinobacteria | 10.78 | 0.000 | 9.88 | 0.000 |
| sensitive | BPA | OTU_202 | Alphaproteobacteria | 5.98 | 0.000 | 4.91 | 0.000 |
| sensitive | BPA | OTU_52 | Planctomycetes | 10.40 | 0.000 | 7.04 | 0.001 |
| sensitive | BPA | OTU_76 | Gammaproteobacteria | 4.67 | 0.000 | 4.30 | 0.000 |
| sensitive | BPA | OTU_117 | Bacteroidetes | 7.65 | 0.000 | 5.22 | 0.002 |
| sensitive | BPA | OTU_65 | Actinobacteria | 11.17 | 0.000 | 11.07 | 0.000 |
| sensitive | BPA | OTU_222 | Bacteroidetes | 9.12 | 0.000 | 8.54 | 0.000 |
| sensitive | BPA | OTU_136 | Planctomycetes | 4.40 | 0.000 | 6.34 | 0.000 |
| sensitive | BPA | OTU_187 | Alphaproteobacteria | 5.00 | 0.000 | 2.95 | 0.013 |
| sensitive | BPA | OTU_131 | Alphaproteobacteria | 4.90 | 0.000 | 3.39 | 0.005 |
| sensitive | BPA | OTU_266 | Bacteroidetes | 6.16 | 0.000 | 7.32 | 0.000 |
| sensitive | BPA | OTU_165 | Bacteroidetes | 9.10 | 0.000 | 5.60 | 0.012 |
| sensitive | BPA | OTU_9 | Alphaproteobacteria | 2.80 | 0.001 | 1.73 | 0.025 |
| sensitive | BPA | OTU_98 | Verrucomicrobia | 3.77 | 0.002 | 4.56 | 0.000 |
| sensitive | BPA | OTU_366 | Bacteroidetes | 8.02 | 0.002 | 7.45 | 0.001 |
| sensitive | BPA | OTU_397 | Alphaproteobacteria | 8.57 | 0.003 | 6.97 | 0.005 |
| sensitive | BPA | OTU_104 | Actinobacteria | 9.34 | 0.004 | 8.77 | 0.001 |

| Ecological category | Treatment | OTUID | Taxonomy (Phylum/class) | log2(P1/P2) | *P.adj* | log2(P2/P3) | *P.adj* |
|---|---|---|---|---|---|---|---|
| sensitive | BPA | OTU_74 | Actinobacteria | 9.71 | 0.005 | 8.81 | 0.003 |
| sensitive | BPA | OTU_113 | Planctomycetes | 3.88 | 0.005 | 6.13 | 0.000 |
| sensitive | BPA | OTU_135 | Alphaproteobacteria | 8.95 | 0.005 | 8.38 | 0.002 |
| sensitive | BPA | OTU_245 | Bacteroidetes | 9.26 | 0.005 | 7.15 | 0.013 |
| sensitive | BPA | OTU_154 | Verrucomicrobia | 5.57 | 0.006 | 4.58 | 0.009 |
| sensitive | BPA | OTU_249 | Alphaproteobacteria | 3.45 | 0.006 | 2.63 | 0.017 |
| sensitive | BPA | OTU_203 | Verrucomicrobia | 7.53 | 0.006 | 6.63 | 0.005 |
| sensitive | BPA | OTU_164 | Gammaproteobacteria | 6.68 | 0.007 | 8.38 | 0.000 |
| sensitive | BPA | OTU_413 | Planctomycetes | 5.20 | 0.010 | 6.03 | 0.000 |
| sensitive | BPA | OTU_137 | Actinobacteria | 9.44 | 0.010 | 8.87 | 0.005 |
| sensitive | BPA | OTU_178 | Verrucomicrobia | 9.18 | 0.011 | 8.61 | 0.005 |
| sensitive | BPA | OTU_268 | Gammaproteobacteria | 3.59 | 0.011 | 4.22 | 0.001 |
| sensitive | BPA | OTU_72 | Gammaproteobacteria | 3.38 | 0.011 | 3.48 | 0.003 |
| sensitive | BPA | OTU_323 | Gammaproteobacteria | 3.53 | 0.013 | 3.15 | 0.011 |
| sensitive | BPA | OTU_462 | Bacteroidetes | 7.97 | 0.014 | 7.39 | 0.008 |
| sensitive | BPA | OTU_90 | Actinobacteria | 8.09 | 0.014 | 7.99 | 0.005 |
| sensitive | BPA | OTU_160 | Actinobacteria | 9.16 | 0.014 | 8.59 | 0.008 |
| sensitive | BPA | OTU_446 | Actinobacteria | 6.23 | 0.015 | 5.65 | 0.010 |
| sensitive | BPA | OTU_357 | Bacteroidetes | 4.74 | 0.020 | 6.07 | 0.001 |
| sensitive | BPA | OTU_335 | Bacteroidetes | 8.78 | 0.020 | 8.21 | 0.012 |
| sensitive | BPA | OTU_170 | Bacteroidetes | 5.27 | 0.024 | 7.43 | 0.000 |
| sensitive | BPA | OTU_256 | Bacteroidetes | 5.25 | 0.025 | 6.07 | 0.003 |
| sensitive | BPA | OTU_278 | Verrucomicrobia | 6.70 | 0.028 | 6.12 | 0.019 |
| sensitive | BPA | OTU_379 | Actinobacteria | 6.93 | 0.029 | 6.36 | 0.020 |
| sensitive | BPA | OTU_195 | Alphaproteobacteria | 3.85 | 0.030 | 4.53 | 0.004 |
| sensitive | BPA | OTU_274 | Verrucomicrobia | 6.44 | 0.030 | 5.86 | 0.022 |
| sensitive | BPA | OTU_166 | Gammaproteobacteria | 3.53 | 0.030 | 3.41 | 0.017 |
| sensitive | BPA | OTU_285 | Cyanobacteria | 5.71 | 0.034 | 4.82 | 0.039 |
| sensitive | BPA | OTU_149 | Planctomycetes | 2.58 | 0.036 | 4.99 | 0.000 |
| sensitive | BPA | OTU_158 | Actinobacteria | 7.93 | 0.036 | 7.35 | 0.025 |
| sensitive | BPA | OTU_234 | Bacteroidetes | 2.94 | 0.040 | 3.50 | 0.005 |
| sensitive | BPA | OTU_313 | Bacteroidetes | 4.60 | 0.040 | 6.97 | 0.001 |
| sensitive | BPA | OTU_439 | Verrucomicrobia | 6.10 | 0.040 | 5.52 | 0.033 |
| sensitive | BPA | OTU_227 | Verrucomicrobia | 7.58 | 0.040 | 7.01 | 0.030 |
| sensitive | BPA | OTU_388 | Gammaproteobacteria | 4.46 | 0.042 | 6.03 | 0.002 |
| sensitive | BPA | OTU_308 | Actinobacteria | 7.58 | 0.044 | 7.00 | 0.035 |
| sensitive | BPA | OTU_470 | Gammaproteobacteria | 4.10 | 0.047 | 4.96 | 0.005 |
| sensitive | BPA | OTU_493 | Planctomycetes | 4.19 | 0.049 | 4.26 | 0.020 |
| sensitive | TCS | OTU_74 | Actinobacteria | 25.10 | 0.000 | 25.31 | 0.000 |
| sensitive | TCS | OTU_110 | Actinobacteria | 25.81 | 0.000 | 26.19 | 0.000 |
| sensitive | TCS | OTU_158 | Actinobacteria | 23.37 | 0.000 | 23.92 | 0.000 |

| Ecological category | Treatment | OTUID | Taxonomy (Phylum/class) | log2(P1/P2) | *P.adj* | log2(P2/P3) | *P.adj* |
|---|---|---|---|---|---|---|---|
| sensitive | TCS | OTU_201 | Verrucomicrobia | 23.15 | 0.000 | 23.85 | 0.000 |
| sensitive | TCS | OTU_27 | Actinobacteria | 11.04 | 0.000 | 11.47 | 0.000 |
| sensitive | TCS | OTU_47 | Actinobacteria | 9.30 | 0.000 | 10.73 | 0.000 |
| sensitive | TCS | OTU_185 | Bacteroidetes | 4.80 | 0.000 | 7.53 | 0.000 |
| sensitive | TCS | OTU_117 | Bacteroidetes | 4.31 | 0.000 | 7.20 | 0.000 |
| sensitive | TCS | OTU_104 | Actinobacteria | 8.59 | 0.001 | 9.57 | 0.000 |
| sensitive | TCS | OTU_165 | Bacteroidetes | 9.55 | 0.001 | 9.56 | 0.000 |
| sensitive | TCS | OTU_72 | Gammaproteobacteria | 8.60 | 0.001 | 8.76 | 0.000 |
| sensitive | TCS | OTU_131 | Alphaproteobacteria | 4.08 | 0.002 | 4.70 | 0.000 |
| sensitive | TCS | OTU_332 | Bacteroidetes | 6.77 | 0.002 | 7.26 | 0.000 |
| sensitive | TCS | OTU_187 | Alphaproteobacteria | 3.77 | 0.002 | 3.41 | 0.001 |
| sensitive | TCS | OTU_81 | Actinobacteria | 8.65 | 0.002 | 9.35 | 0.000 |
| sensitive | TCS | OTU_195 | Alphaproteobacteria | 4.34 | 0.002 | 3.65 | 0.001 |
| sensitive | TCS | OTU_222 | Bacteroidetes | 6.33 | 0.002 | 6.34 | 0.000 |
| sensitive | TCS | OTU_90 | Actinobacteria | 9.46 | 0.002 | 9.63 | 0.000 |
| sensitive | TCS | OTU_180 | Gammaproteobacteria | 6.21 | 0.005 | 4.03 | 0.019 |
| sensitive | TCS | OTU_359 | Alphaproteobacteria | 6.84 | 0.005 | 7.47 | 0.000 |
| sensitive | TCS | OTU_65 | Actinobacteria | 7.92 | 0.006 | 8.41 | 0.000 |
| sensitive | TCS | OTU_162 | Bacteroidetes | 5.24 | 0.006 | 4.52 | 0.004 |
| sensitive | TCS | OTU_130 | Alphaproteobacteria | 2.65 | 0.007 | 2.13 | 0.010 |
| sensitive | TCS | OTU_178 | Verrucomicrobia | 9.03 | 0.009 | 9.04 | 0.001 |
| sensitive | TCS | OTU_136 | Planctomycetes | 5.47 | 0.010 | 9.05 | 0.000 |
| sensitive | TCS | OTU_412 | Alphaproteobacteria | 7.18 | 0.010 | 7.67 | 0.001 |
| sensitive | TCS | OTU_164 | Gammaproteobacteria | 5.18 | 0.010 | 7.31 | 0.000 |
| sensitive | TCS | OTU_135 | Alphaproteobacteria | 6.66 | 0.014 | 6.67 | 0.003 |
| sensitive | TCS | OTU_245 | Bacteroidetes | 8.32 | 0.014 | 8.32 | 0.003 |
| sensitive | TCS | OTU_170 | Bacteroidetes | 5.25 | 0.016 | 6.52 | 0.000 |
| sensitive | TCS | OTU_76 | Gammaproteobacteria | 2.66 | 0.016 | 4.20 | 0.000 |
| sensitive | TCS | OTU_313 | Bacteroidetes | 5.18 | 0.016 | 7.15 | 0.000 |
| sensitive | TCS | OTU_160 | Actinobacteria | 8.98 | 0.022 | 8.99 | 0.005 |
| sensitive | TCS | OTU_235 | Planctomycetes | 5.64 | 0.023 | 6.61 | 0.001 |
| sensitive | TCS | OTU_395 | Bacteroidetes | 5.37 | 0.023 | 5.35 | 0.006 |
| sensitive | TCS | OTU_137 | Actinobacteria | 8.62 | 0.025 | 8.63 | 0.006 |
| sensitive | TCS | OTU_203 | Verrucomicrobia | 7.31 | 0.028 | 7.32 | 0.007 |
| sensitive | TCS | OTU_262 | Alphaproteobacteria | 4.76 | 0.033 | 4.14 | 0.021 |
| sensitive | TCS | OTU_218 | Gammaproteobacteria | 5.10 | 0.034 | 6.51 | 0.001 |
| sensitive | TCS | OTU_190 | Bacteroidetes | 4.74 | 0.035 | 3.84 | 0.036 |
| sensitive | TCS | OTU_433 | Alphaproteobacteria | 4.12 | 0.035 | 3.68 | 0.019 |
| sensitive | TCS | OTU_60 | Actinobacteria | 8.29 | 0.035 | 8.30 | 0.010 |
| sensitive | TCS | OTU_454 | Gammaproteobacteria | 3.75 | 0.036 | 3.39 | 0.017 |
| sensitive | TCS | OTU_365 | Actinobacteria | 8.49 | 0.036 | 8.50 | 0.011 |

| Ecological category | Treatment | OTUID | Taxonomy (Phylum/class) | log2(P1/P2) | *P.adj* | log2(P2/P3) | *P.adj* |
|---|---|---|---|---|---|---|---|
| sensitive | TCS | OTU_278 | Verrucomicrobia | 6.89 | 0.036 | 6.90 | 0.010 |
| sensitive | TCS | OTU_276 | Bacteroidetes | 5.49 | 0.041 | 7.77 | 0.001 |
| sensitive | TCS | OTU_16 | Planctomycetes | 3.19 | 0.047 | 4.45 | 0.001 |
| sensitive | MI | OTU_274 | Verrucomicrobia | 23.30 | 0.000 | 23.62 | 0.000 |
| sensitive | MI | OTU_19 | Planctomycetes | 9.26 | 0.000 | 8.43 | 0.000 |
| sensitive | MI | OTU_136 | Planctomycetes | 8.20 | 0.000 | 9.73 | 0.000 |
| sensitive | MI | OTU_47 | Actinobacteria | 9.77 | 0.000 | 10.88 | 0.000 |
| sensitive | MI | OTU_27 | Actinobacteria | 11.48 | 0.000 | 11.68 | 0.000 |
| sensitive | MI | OTU_117 | Bacteroidetes | 7.62 | 0.000 | 4.86 | 0.001 |
| sensitive | MI | OTU_3 | Planctomycetes | 5.89 | 0.000 | 4.61 | 0.001 |
| sensitive | MI | OTU_60 | Actinobacteria | 10.37 | 0.001 | 10.09 | 0.000 |
| sensitive | MI | OTU_104 | Actinobacteria | 8.47 | 0.001 | 8.67 | 0.000 |
| sensitive | MI | OTU_65 | Actinobacteria | 10.49 | 0.002 | 10.21 | 0.000 |
| sensitive | MI | OTU_76 | Gammaproteobacteria | 4.09 | 0.002 | 6.00 | 0.000 |
| sensitive | MI | OTU_72 | Gammaproteobacteria | 7.17 | 0.002 | 8.00 | 0.000 |
| sensitive | MI | OTU_164 | Gammaproteobacteria | 3.96 | 0.002 | 5.70 | 0.000 |
| sensitive | MI | OTU_222 | Bacteroidetes | 6.72 | 0.002 | 5.88 | 0.002 |
| sensitive | MI | OTU_81 | Actinobacteria | 9.49 | 0.003 | 9.21 | 0.001 |
| sensitive | MI | OTU_332 | Bacteroidetes | 5.97 | 0.004 | 5.04 | 0.005 |
| sensitive | MI | OTU_366 | Bacteroidetes | 7.12 | 0.004 | 7.32 | 0.001 |
| sensitive | MI | OTU_131 | Alphaproteobacteria | 3.87 | 0.005 | 4.55 | 0.000 |
| sensitive | MI | OTU_74 | Actinobacteria | 9.44 | 0.005 | 9.64 | 0.001 |
| sensitive | MI | OTU_16 | Planctomycetes | 5.70 | 0.005 | 5.47 | 0.002 |
| sensitive | MI | OTU_235 | Planctomycetes | 7.94 | 0.005 | 8.62 | 0.000 |
| sensitive | MI | OTU_178 | Verrucomicrobia | 7.81 | 0.007 | 7.53 | 0.002 |
| sensitive | MI | OTU_113 | Planctomycetes | 3.49 | 0.009 | 9.68 | 0.000 |
| sensitive | MI | OTU_90 | Actinobacteria | 8.19 | 0.009 | 8.07 | 0.003 |
| sensitive | MI | OTU_433 | Alphaproteobacteria | 5.31 | 0.009 | 6.11 | 0.001 |
| sensitive | MI | OTU_467 | Bacteroidetes | 6.40 | 0.009 | 4.22 | 0.044 |
| sensitive | MI | OTU_170 | Bacteroidetes | 5.25 | 0.009 | 7.54 | 0.000 |
| sensitive | MI | OTU_308 | Actinobacteria | 7.75 | 0.011 | 7.47 | 0.005 |
| sensitive | MI | OTU_107 | Gammaproteobacteria | 2.20 | 0.015 | 2.20 | 0.005 |
| sensitive | MI | OTU_165 | Bacteroidetes | 7.28 | 0.015 | 8.45 | 0.001 |
| sensitive | MI | OTU_31 | Bacteroidetes | 4.53 | 0.015 | 6.03 | 0.000 |
| sensitive | MI | OTU_184 | Planctomycetes | 3.27 | 0.015 | 8.79 | 0.000 |
| sensitive | MI | OTU_200 | Planctomycetes | 4.49 | 0.015 | 8.62 | 0.000 |
| sensitive | MI | OTU_404 | Gammaproteobacteria | 4.56 | 0.017 | 5.37 | 0.001 |
| sensitive | MI | OTU_195 | Alphaproteobacteria | 3.62 | 0.021 | 3.88 | 0.004 |
| sensitive | MI | OTU_9 | Alphaproteobacteria | 2.12 | 0.022 | 1.84 | 0.023 |
| sensitive | MI | OTU_130 | Alphaproteobacteria | 2.43 | 0.022 | 2.64 | 0.004 |
| sensitive | MI | OTU_7 | Bacteroidetes | 3.45 | 0.023 | 4.21 | 0.001 |

| Ecological category | Treatment | OTUID | Taxonomy (Phylum/class) | log2(P1/P2) | *P.adj* | log2(P2/P3) | *P.adj* |
|---|---|---|---|---|---|---|---|
| sensitive | MI | OTU_75 | Bacteroidetes | 6.34 | 0.024 | 4.84 | 0.044 |
| sensitive | MI | OTU_239 | Bacteroidetes | 4.27 | 0.024 | 3.53 | 0.032 |
| sensitive | MI | OTU_135 | Alphaproteobacteria | 6.44 | 0.026 | 7.12 | 0.004 |
| sensitive | MI | OTU_35 | Bacteroidetes | 3.48 | 0.027 | 3.74 | 0.006 |
| sensitive | MI | OTU_285 | Cyanobacteria | 8.28 | 0.027 | 8.00 | 0.014 |
| sensitive | MI | OTU_304 | Verrucomicrobia | 7.15 | 0.027 | 7.03 | 0.012 |
| sensitive | MI | OTU_244 | Actinobacteria | 6.89 | 0.030 | 6.61 | 0.016 |
| sensitive | MI | OTU_204 | Gammaproteobacteria | 3.07 | 0.031 | 3.05 | 0.014 |
| sensitive | MI | OTU_169 | Actinobacteria | 8.08 | 0.036 | 7.80 | 0.019 |
| sensitive | MI | OTU_137 | Actinobacteria | 7.94 | 0.038 | 8.15 | 0.013 |
| sensitive | MI | OTU_160 | Actinobacteria | 7.93 | 0.038 | 8.13 | 0.013 |
| sensitive | MI | OTU_124 | Bacteroidetes | 3.65 | 0.045 | 6.16 | 0.000 |
| sensitive | MI | OTU_496 | Actinobacteria | 6.46 | 0.048 | 6.18 | 0.027 |
| sensitive | MII | OTU_16 | Planctomycetes | 9.52 | 0.000 | 10.53 | 0.000 |
| sensitive | MII | OTU_3 | Planctomycetes | 8.01 | 0.000 | 7.54 | 0.000 |
| sensitive | MII | OTU_19 | Planctomycetes | 8.06 | 0.000 | 9.42 | 0.000 |
| sensitive | MII | OTU_131 | Alphaproteobacteria | 6.62 | 0.000 | 4.19 | 0.002 |
| sensitive | MII | OTU_195 | Alphaproteobacteria | 7.37 | 0.000 | 4.83 | 0.001 |
| sensitive | MII | OTU_27 | Actinobacteria | 10.70 | 0.000 | 10.60 | 0.000 |
| sensitive | MII | OTU_47 | Actinobacteria | 8.81 | 0.000 | 10.00 | 0.000 |
| sensitive | MII | OTU_187 | Alphaproteobacteria | 6.15 | 0.000 | 3.74 | 0.004 |
| sensitive | MII | OTU_113 | Planctomycetes | 5.79 | 0.000 | 9.58 | 0.000 |
| sensitive | MII | OTU_104 | Actinobacteria | 9.14 | 0.001 | 8.57 | 0.000 |
| sensitive | MII | OTU_165 | Bacteroidetes | 9.51 | 0.001 | 8.94 | 0.000 |
| sensitive | MII | OTU_52 | Planctomycetes | 6.97 | 0.001 | 8.40 | 0.000 |
| sensitive | MII | OTU_180 | Gammaproteobacteria | 6.11 | 0.001 | 4.04 | 0.008 |
| sensitive | MII | OTU_72 | Gammaproteobacteria | 8.76 | 0.001 | 9.76 | 0.000 |
| sensitive | MII | OTU_166 | Gammaproteobacteria | 7.60 | 0.002 | 7.81 | 0.000 |
| sensitive | MII | OTU_136 | Planctomycetes | 5.85 | 0.002 | 8.73 | 0.000 |
| sensitive | MII | OTU_154 | Verrucomicrobia | 4.00 | 0.002 | 7.90 | 0.000 |
| sensitive | MII | OTU_347 | Gammaproteobacteria | 8.28 | 0.003 | 7.86 | 0.001 |
| sensitive | MII | OTU_74 | Actinobacteria | 9.35 | 0.005 | 8.77 | 0.002 |
| sensitive | MII | OTU_262 | Alphaproteobacteria | 5.42 | 0.006 | 4.23 | 0.011 |
| sensitive | MII | OTU_76 | Gammaproteobacteria | 4.77 | 0.006 | 3.81 | 0.011 |
| sensitive | MII | OTU_90 | Actinobacteria | 9.37 | 0.008 | 9.28 | 0.002 |
| sensitive | MII | OTU_257 | Planctomycetes | 5.76 | 0.009 | 7.19 | 0.000 |
| sensitive | MII | OTU_65 | Actinobacteria | 8.08 | 0.009 | 7.98 | 0.002 |
| sensitive | MII | OTU_222 | Bacteroidetes | 7.84 | 0.010 | 7.26 | 0.004 |
| sensitive | MII | OTU_460 | Verrucomicrobia | 6.77 | 0.011 | 6.19 | 0.005 |
| sensitive | MII | OTU_164 | Gammaproteobacteria | 5.28 | 0.011 | 7.20 | 0.000 |
| sensitive | MII | OTU_81 | Actinobacteria | 8.50 | 0.012 | 8.40 | 0.003 |

| Ecological category | Treatment | OTUID | Taxonomy (Phylum/class) | log2(P1/P2) | *P.adj* | log2(P2/P3) | *P.adj* |
|---|---|---|---|---|---|---|---|
| sensitive | MII | OTU_448 | Planctomycetes | 6.56 | 0.012 | 5.98 | 0.007 |
| sensitive | MII | OTU_137 | Actinobacteria | 7.23 | 0.013 | 7.14 | 0.004 |
| sensitive | MII | OTU_197 | Alphaproteobacteria | 8.42 | 0.013 | 8.32 | 0.004 |
| sensitive | MII | OTU_357 | Bacteroidetes | 5.37 | 0.016 | 5.45 | 0.003 |
| sensitive | MII | OTU_235 | Planctomycetes | 6.12 | 0.016 | 7.99 | 0.000 |
| sensitive | MII | OTU_135 | Alphaproteobacteria | 9.22 | 0.017 | 8.65 | 0.008 |
| sensitive | MII | OTU_470 | Gammaproteobacteria | 4.41 | 0.020 | 4.91 | 0.002 |
| sensitive | MII | OTU_60 | Actinobacteria | 8.90 | 0.027 | 8.33 | 0.015 |
| sensitive | MII | OTU_358 | Bacteroidetes | 7.19 | 0.030 | 6.62 | 0.018 |
| sensitive | MII | OTU_379 | Actinobacteria | 6.82 | 0.039 | 6.25 | 0.023 |
| sensitive | MII | OTU_178 | Verrucomicrobia | 7.23 | 0.041 | 7.63 | 0.008 |
| sensitive | MII | OTU_160 | Actinobacteria | 7.82 | 0.041 | 7.25 | 0.023 |
| sensitive | MII | OTU_224 | Actinobacteria | 8.14 | 0.043 | 7.57 | 0.024 |
| tolerant | BPA | OTU_109 | Alphaproteobacteria | -6.09 | 0.000 | -3.38 | 0.002 |
| tolerant | BPA | OTU_48 | Alphaproteobacteria | -10.09 | 0.000 | -11.95 | 0.000 |
| tolerant | BPA | OTU_78 | Gammaproteobacteria | -6.09 | 0.000 | -8.93 | 0.000 |
| tolerant | BPA | OTU_14 | Verrucomicrobia | -7.57 | 0.000 | -6.63 | 0.003 |
| tolerant | BPA | OTU_186 | Bacteroidetes | -6.82 | 0.005 | -5.89 | 0.047 |
| tolerant | BPA | OTU_134 | Bacteroidetes | -3.59 | 0.010 | -4.81 | 0.003 |
| tolerant | TCS | OTU_48 | Alphaproteobacteria | -10.34 | 0.000 | -10.30 | 0.000 |
| tolerant | TCS | OTU_78 | Gammaproteobacteria | -7.98 | 0.000 | -8.42 | 0.000 |
| tolerant | TCS | OTU_215 | Bacteroidetes | -9.44 | 0.000 | -4.81 | 0.022 |
| tolerant | TCS | OTU_273 | Bacteroidetes | -8.76 | 0.000 | -6.79 | 0.004 |
| tolerant | TCS | OTU_250 | Verrucomicrobia | -5.90 | 0.000 | -3.85 | 0.047 |
| tolerant | TCS | OTU_306 | Gammaproteobacteria | -7.14 | 0.000 | -7.11 | 0.004 |
| tolerant | TCS | OTU_372 | Gammaproteobacteria | -7.10 | 0.001 | -6.10 | 0.013 |
| tolerant | TCS | OTU_75 | Bacteroidetes | -4.05 | 0.002 | -4.91 | 0.002 |
| tolerant | TCS | OTU_102 | Bacteroidetes | -3.94 | 0.002 | -6.17 | 0.000 |
| tolerant | TCS | OTU_307 | Planctomycetes | -3.09 | 0.004 | -2.69 | 0.037 |
| tolerant | TCS | OTU_33 | Alphaproteobacteria | -3.30 | 0.004 | -3.77 | 0.005 |
| tolerant | TCS | OTU_138 | Gammaproteobacteria | -3.27 | 0.015 | -3.31 | 0.047 |
| tolerant | TCS | OTU_134 | Bacteroidetes | -3.05 | 0.015 | -3.43 | 0.022 |
| tolerant | TCS | OTU_216 | Gammaproteobacteria | -4.57 | 0.028 | -5.87 | 0.022 |
| tolerant | TCS | OTU_279 | Gammaproteobacteria | -6.72 | 0.029 | -22.59 | 0.000 |
| tolerant | MI | OTU_78 | Gammaproteobacteria | -9.44 | 0.000 | -8.99 | 0.000 |
| tolerant | MI | OTU_48 | Alphaproteobacteria | -11.47 | 0.000 | -11.48 | 0.000 |
| tolerant | MI | OTU_33 | Alphaproteobacteria | -4.84 | 0.000 | -3.60 | 0.026 |
| tolerant | MI | OTU_116 | Chlamydiae | -8.16 | 0.001 | -7.39 | 0.005 |
| tolerant | MI | OTU_252 | Bacteroidetes | -7.95 | 0.001 | -5.38 | 0.047 |
| tolerant | MI | OTU_128 | Gammaproteobacteria | -7.43 | 0.002 | -7.98 | 0.004 |
| tolerant | MI | OTU_156 | Bacteroidetes | -4.16 | 0.010 | -4.01 | 0.031 |

| Ecological category | Treatment | OTUID | Taxonomy (Phylum/class) | log2(P1/P2) | *P.adj* | log2(P2/P3) | *P.adj* |
|---|---|---|---|---|---|---|---|
| tolerant | MI | OTU_146 | Verrucomicrobia | -6.91 | 0.014 | -7.40 | 0.025 |
| tolerant | MI | OTU_253 | Bacteroidetes | -5.74 | 0.017 | -6.17 | 0.027 |
| tolerant | MI | OTU_49 | Bacteroidetes | -5.05 | 0.017 | -5.25 | 0.031 |
| tolerant | MI | OTU_306 | Gammaproteobacteria | -5.32 | 0.020 | -5.81 | 0.029 |
| tolerant | MI | OTU_490 | Verrucomicrobia | -5.62 | 0.021 | -6.11 | 0.031 |
| tolerant | MII | OTU_211 | Bacteroidetes | -22.86 | 0.000 | -7.68 | 0.010 |
| tolerant | MII | OTU_44 | Alphaproteobacteria | -24.61 | 0.000 | -9.16 | 0.014 |
| tolerant | MII | OTU_151 | Alphaproteobacteria | -23.63 | 0.000 | -27.79 | 0.000 |
| tolerant | MII | OTU_78 | Gammaproteobacteria | -8.44 | 0.000 | -8.25 | 0.000 |
| tolerant | MII | OTU_58 | Gammaproteobacteria | -6.53 | 0.000 | -3.45 | 0.024 |
| tolerant | MII | OTU_48 | Alphaproteobacteria | -10.14 | 0.000 | -10.51 | 0.000 |
| tolerant | MII | OTU_134 | Bacteroidetes | -7.74 | 0.000 | -5.92 | 0.000 |
| tolerant | MII | OTU_306 | Gammaproteobacteria | -6.50 | 0.000 | -6.40 | 0.000 |
| tolerant | MII | OTU_77 | Alphaproteobacteria | -9.15 | 0.001 | -8.55 | 0.010 |
| tolerant | MII | OTU_372 | Gammaproteobacteria | -6.10 | 0.009 | -6.48 | 0.022 |

**Table S4** The list of ecologically-grouped OTUs (opportunistic, sensitive and tolerant) shared across treatments. Max_rel: maximal relative abundance of the OTUs in all samples; max_id: the sample ID in which the maximal relative abundance was detected, e.g., denoted by "batch_treatment_replicate", sample 'B1_CON_002' represents the number of batch, treatment id and the number of the biological replicate in that treatment. Treatment IDs are: 'Initial' containing the starting bacterial communities before the experimental implementation; 'Con' microcosms containing no micropollutant additions and serve as the controls; 'BPA' microcosms containing bisphenol A; 'TCS' microcosms containing triclosan; 'MI' microcosms containing a mixture of bisphenol A and triclosan; 'MII' microcosms containing a mixture of bisphenol A, triclosan, bisphenol-S and triclocarban.

| OTUID | Treatment | | | | Taxonomic affiliation | | | | | max_rel | max_id |
|---|---|---|---|---|---|---|---|---|---|---|---|
| | BPA | TCS | MI | MII | Phylum | Class | Order | Family | Genus | | |
| OTU_104 | sensitive | sensitive | sensitive | sensitive | Actinobacteria | Actinobacteria | Frankiales | Sporichthyaceae | hgcI clade | 6.34% | Initial_002 |
| OTU_137 | sensitive | sensitive | sensitive | sensitive | Actinobacteria | Actinobacteria | Frankiales | Sporichthyaceae | Ambiguous_taxa | 1.66% | Initial_002 |
| OTU_158 | sensitive | sensitive | | | Actinobacteria | Actinobacteria | Frankiales | Sporichthyaceae | Candidatus Planktophila | 4.29% | Initial_002 |
| OTU_160 | sensitive | sensitive | sensitive | sensitive | Actinobacteria | Actinobacteria | Frankiales | Sporichthyaceae | hgcI clade | 0.97% | Initial_001 |
| OTU_169 | sensitive | | sensitive | | Actinobacteria | Actinobacteria | Frankiales | Sporichthyaceae | hgcI clade | 3.86% | B1_BPA_001 |
| OTU_27 | sensitive | sensitive | sensitive | sensitive | Actinobacteria | Actinobacteria | Frankiales | Sporichthyaceae | hgcI clade | 15.94% | Initial_002 |
| OTU_308 | sensitive | | sensitive | | Actinobacteria | Actinobacteria | Corynebacteriales | Mycobacteriaceae | Mycobacterium | 0.28% | B1_BPA_001 |
| OTU_379 | sensitive | | | sensitive | Actinobacteria | Thermoleophilia | Solirubrobacterales | Solirubrobacteraceae | Conexibacter | 0.16% | B1_MII_001 |
| OTU_47 | sensitive | sensitive | sensitive | sensitive | Actinobacteria | Actinobacteria | Frankiales | Sporichthyaceae | hgcI clade | 8.32% | Initial_002 |
| OTU_60 | sensitive | sensitive | sensitive | sensitive | Actinobacteria | Acidimicrobiia | Microtrichales | Ilumatobacteraceae | CL500-29 marine group | 7.72% | Initial_003 |
| OTU_65 | sensitive | sensitive | sensitive | sensitive | Actinobacteria | Actinobacteria | Frankiales | Sporichthyaceae | hgcI clade | 11.65% | Initial_001 |
| OTU_74 | sensitive | sensitive | sensitive | sensitive | Actinobacteria | Actinobacteria | Frankiales | Sporichthyaceae | hgcI clade | 13.28% | Initial_002 |
| OTU_81 | sensitive | sensitive | sensitive | sensitive | Actinobacteria | Actinobacteria | Frankiales | Sporichthyaceae | | 4.46% | Initial_002 |
| OTU_90 | sensitive | sensitive | sensitive | sensitive | Actinobacteria | Acidimicrobiia | Microtrichales | Ilumatobacteraceae | CL500-29 marine group | 5.01% | Initial_002 |
| OTU_130 | | sensitive | sensitive | | Proteobacteria | Alphaproteobacteria | Rhizobiales | Beijerinckiaceae | Methylobacterium | 0.56% | Initial_002 |
| OTU_131 | sensitive | sensitive | sensitive | sensitive | Proteobacteria | Alphaproteobacteria | Rhizobiales | Beijerinckiaceae | Methylobacterium | 0.86% | B1_MII_003 |
| OTU_135 | sensitive | sensitive | sensitive | sensitive | Proteobacteria | Alphaproteobacteria | SAR11 clade | Clade III | uncultured bacterium | 2.60% | Initial_001 |
| OTU_187 | sensitive | sensitive | | sensitive | Proteobacteria | Alphaproteobacteria | Caulobacterales | Caulobacteraceae | Caulobacter | 0.40% | B1_MII_003 |
| OTU_195 | sensitive | sensitive | sensitive | sensitive | Proteobacteria | Alphaproteobacteria | Rhizobiales | Beijerinckiaceae | Methylobacterium | 0.76% | B1_MII_003 |
| OTU_33 | | tolerant | tolerant | | Proteobacteria | Alphaproteobacteria | Sphingomonadales | Sphingomonadaceae | Novosphingobium | 30.48% | B6_BPA_001 |
| OTU_433 | | sensitive | sensitive | | Proteobacteria | Alphaproteobacteria | Rhizobiales | Beijerinckiaceae | Methylobacterium | 0.14% | B1_MI_003 |
| OTU_48 | tolerant | tolerant | tolerant | tolerant | Proteobacteria | Alphaproteobacteria | Sphingomonadales | Sphingomonadaceae | Sphingomonas | 9.71% | B7_MI_001 |

| OTUID | Treatment | | | | Taxonomic affiliation | | | | | max_rel | max_id |
|---|---|---|---|---|---|---|---|---|---|---|---|
| | BPA | TCS | MI | MII | Phylum | Class | Order | Family | Genus | | |
| OTU_9 | sensitive | | sensitive | | Proteobacteria | Alphaproteobacteria | Rhizobiales | Rhizobiaceae | Shinella | 9.26% | B1_MII_002 |
| OTU_107 | sensitive | | sensitive | | Proteobacteria | Gammaproteobacteria | Betaproteobacteriales | Rhodocyclaceae | Methyloversatilis | 0.66% | B4_TCS_001 |
| OTU_164 | sensitive | sensitive | sensitive | sensitive | Proteobacteria | Gammaproteobacteria | Betaproteobacteriales | Burkholderiaceae | MWH-UniP1 aquatic group | 0.83% | B1_MII_002 |
| OTU_166 | sensitive | | | sensitive | Proteobacteria | Gammaproteobacteria | Betaproteobacteriales | Burkholderiaceae | Polynucleobacter | 3.42% | B1_MII_001 |
| OTU_180 | | sensitive | | sensitive | Proteobacteria | Gammaproteobacteria | Betaproteobacteriales | Methylophilaceae | Methylotenera | 0.43% | B1_CON_002 |
| OTU_470 | sensitive | | | sensitive | Proteobacteria | Gammaproteobacteria | Betaproteobacteriales | Nitrosomonadaceae | Ellin6067 | 0.04% | B1_BPA_001 |
| OTU_72 | sensitive | sensitive | sensitive | sensitive | Proteobacteria | Gammaproteobacteria | Betaproteobacteriales | Burkholderiaceae | | 2.71% | B1_BPA_002 |
| OTU_76 | sensitive | sensitive | sensitive | sensitive | Proteobacteria | Gammaproteobacteria | Betaproteobacteriales | Burkholderiaceae | Polynucleobacter | 4.13% | B1_MII_001 |
| OTU_78 | tolerant | tolerant | tolerant | tolerant | Proteobacteria | Gammaproteobacteria | Betaproteobacteriales | Burkholderiaceae | Hydrogenophaga | 4.26% | B6_CON_003 |
| OTU_306 | | tolerant | tolerant | tolerant | Proteobacteria | Gammaproteobacteria | Xanthomonadales | Xanthomonadaceae | Silanimonas | 0.14% | B5_MI_003 |
| OTU_372 | | tolerant | | tolerant | Proteobacteria | Gammaproteobacteria | Xanthomonadales | Xanthomonadaceae | Silanimonas | 0.17% | B5_TCS_003 |
| OTU_117 | sensitive | sensitive | sensitive | | Bacteroidetes | Bacteroidia | Flavobacteriales | Flavobacteriaceae | Flavobacterium | 2.66% | B1_CON_001 |
| OTU_134 | tolerant | tolerant | | tolerant | Bacteroidetes | Bacteroidia | Flavobacteriales | Flavobacteriaceae | Flavobacterium | 1.24% | B6_MII_001 |
| OTU_165 | sensitive | sensitive | sensitive | sensitive | Bacteroidetes | Bacteroidia | Sphingobacteriales | env.OPS 17 | | 1.01% | B1_BPA_002 |
| OTU_170 | sensitive | sensitive | sensitive | | Bacteroidetes | Bacteroidia | Chitinophagales | Chitinophagaceae | | 1.43% | B6_MI_002 |
| OTU_185 | sensitive | sensitive | | | Bacteroidetes | Bacteroidia | Sphingobacteriales | | | 0.33% | B2_BPA_001 |
| OTU_222 | sensitive | sensitive | sensitive | sensitive | Bacteroidetes | Bacteroidia | Chitinophagales | 37-13 | | 1.18% | B2_BPA_001 |
| OTU_245 | sensitive | sensitive | | | Bacteroidetes | Bacteroidia | Sphingobacteriales | env.OPS 17 | | 1.07% | B1_BPA_002 |
| OTU_313 | sensitive | sensitive | | | Bacteroidetes | Bacteroidia | Chitinophagales | Chitinophagaceae | | 0.34% | B1_MI_002 |
| OTU_332 | | sensitive | sensitive | | Bacteroidetes | Bacteroidia | Sphingobacteriales | | | 0.17% | B1_TCS_001 |
| OTU_357 | sensitive | | | sensitive | Bacteroidetes | Bacteroidia | Flavobacteriales | Flavobacteriaceae | Flavobacterium | 0.31% | B1_MI_002 |
| OTU_366 | sensitive | | sensitive | | Bacteroidetes | Bacteroidia | Chitinophagales | Chitinophagaceae | Heliimonas | 0.35% | B1_BPA_003 |
| OTU_75 | | tolerant | sensitive | | Bacteroidetes | Bacteroidia | Chitinophagales | Chitinophagaceae | | 11.06% | B6_TCS_002 |

| OTUID | Treatment | | | | Taxonomic affiliation | | | | | max_rel | max_id |
|---|---|---|---|---|---|---|---|---|---|---|---|
| | BPA | TCS | MI | MII | Phylum | Class | Order | Family | Genus | | |
| OTU_285 | sensitive | | sensitive | | Cyanobacteria | Oxyphotobacteria | Synechococcales | Cyanobiaceae | Cyanobium PCC-6307 | 0.38% | Initial_001 |
| OTU_113 | sensitive | | sensitive | sensitive | Planctomycetes | Planctomycetacia | Planctomycetales | Schlesneriaceae | Planctopirus | 1.41% | B1_MII_003 |
| OTU_136 | sensitive | sensitive | sensitive | sensitive | Planctomycetes | Planctomycetacia | Pirellulales | Pirellulaceae | uncultured | 0.95% | B1_MII_001 |
| OTU_16 | | sensitive | sensitive | sensitive | Planctomycetes | Planctomycetacia | Pirellulales | Pirellulaceae | Pirellula | 16.49% | B1_MII_003 |
| OTU_19 | sensitive | | sensitive | sensitive | Planctomycetes | Planctomycetacia | Pirellulales | Pirellulaceae | Pirellula | 9.59% | B2_MI_002 |
| OTU_235 | | sensitive | sensitive | sensitive | Planctomycetes | Planctomycetacia | Pirellulales | Pirellulaceae | uncultured | 0.49% | B1_CON_003 |
| OTU_3 | sensitive | | sensitive | sensitive | Planctomycetes | Planctomycetacia | Pirellulales | Pirellulaceae | Pirellula | 27.65% | B2_CON_003 |
| OTU_52 | sensitive | | | sensitive | Planctomycetes | Planctomycetacia | Pirellulales | Pirellulaceae | Pirellula | 5.42% | B3_TCS_001 |
| OTU_154 | sensitive | | | sensitive | Verrucomicrobia | Verrucomicrobiae | Verrucomicrobiales | Verrucomicrobiaceae | uncultured | 0.56% | B2_CON_003 |
| OTU_178 | sensitive | sensitive | sensitive | sensitive | Verrucomicrobia | Verrucomicrobiae | Methylacidiphilales | Methylacidiphilaceae | uncultured | 0.95% | B1_TCS_002 |
| OTU_203 | sensitive | sensitive | | | Verrucomicrobia | Verrucomicrobiae | Methylacidiphilales | Methylacidiphilaceae | uncultured | 1.17% | Initial_002 |
| OTU_274 | sensitive | | sensitive | | Verrucomicrobia | Verrucomicrobiae | Methylacidiphilales | Methylacidiphilaceae | uncultured | 0.56% | B1_MI_002 |
| OTU_278 | sensitive | sensitive | | | Verrucomicrobia | Verrucomicrobiae | Methylacidiphilales | Methylacidiphilaceae | uncultured | 0.57% | Initial_001 |

**Figure S1**. The location of the Shidou Reservoir in the map of Xiamen city (A) and sampling site in the reservoir (B).

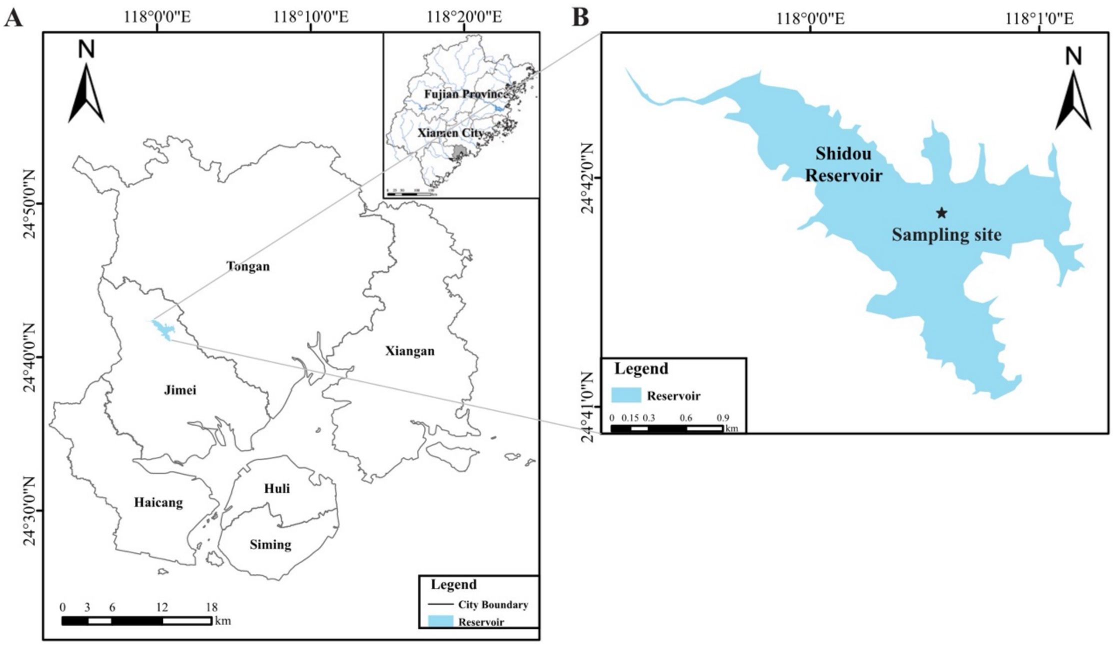

**Figure S2** The removal rates of the micropollutants in sterile microcosms spiked with1 μg/L bisphenol A (BPA), 0.1 μg/L bisphenol S (BPS), 1 μg/L triclosan (TCS) and 0.1 μg/L triclocarban (TCC) for 7-day incubation. The removal rates were presented as the average of the triplicates.

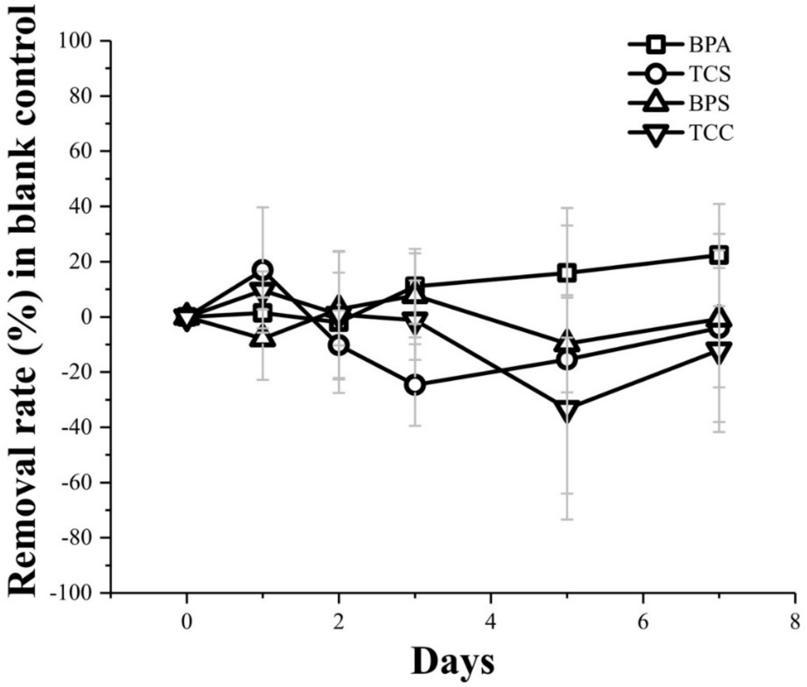

**Figure S3** The variations in concentration of 1 mg/L bisphenol A (BPA), 1 mg/L triclosan (TCS) and 1 mg/L mixed BPA+TCS under the bio-transformation of prokaryotic cultures before (a) and after (b) microcosm cultivation at 1 μg/L level. The prokaryotic cultures used in 1 mg/L bio-transformation experiments were transferred from 1 μg/L microcosm group BPA, TCS and MI respectively. Curves in black and red represent the concentration variations of BPA and TCS in the respective treatment, respectively.

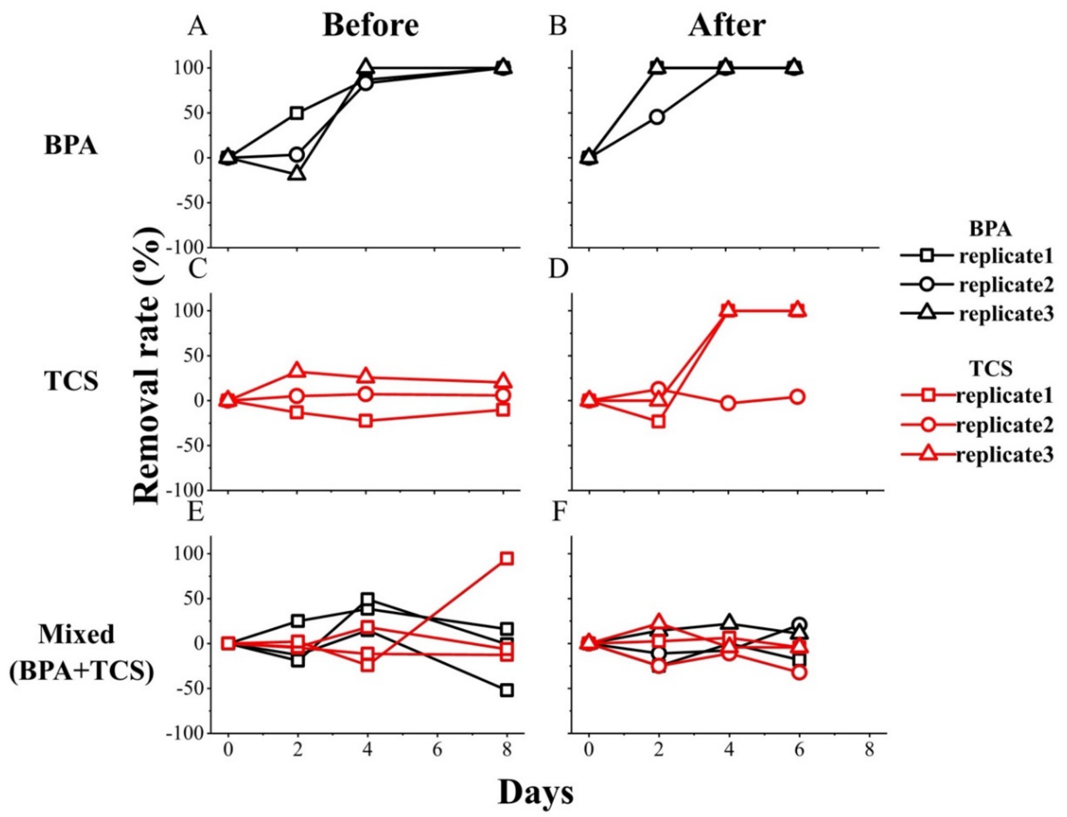

**Figure S4**. The temporal variation of abundance-weighted average 16S rRNA gene copy numbers of bacterial communities in the controls and micropollutant-treated microcosms (A-E), of Pearson's correlation for pairwise treatment comparison (F) and the grouping of similar frequency of gene copy numbers (G). Color gradients in (F) represent Pearson's correlation coefficients. A hierarchical dendrogram (G) illustrated the similar frequency (based on Bray-Curtis dissimilarity) of 16S rRNA gene copy numbers of the communities in the micropollutants-treated batches. Three growth progression phrases across all microcosm batches were identified based on similar average gene copy numbers and the time continuity of the inoculation phase 1 (B1–B2), phase 2 (B3–B4), and phase 3 (B5–B7).

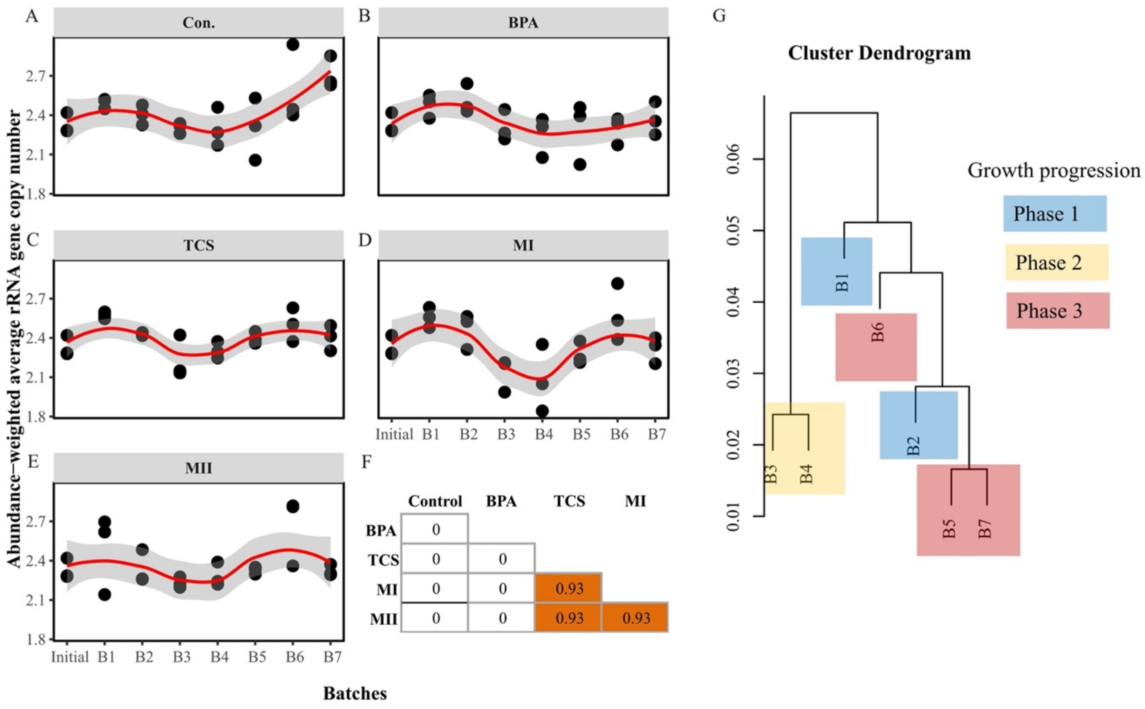

**Figure S5** Percent removal of bisphenol A (BPA), triclosan (TCS), bisphenol S (BPS), and triclocarbon (TCC) from the microcosms. (A) Removal in the single-micropollutant (BPA and TCA) treatments; (B) removal in the mixed-micropollutant (MI and MII) treatments. The open circles represent the removal percentage obtained from each microcosm batch. Tukey's *post-hoc* test was used to determine at which micropollutant the differences in rates occurred, and in this case the removal rate of the micropollutants from each microcosm batch was used as replicates for the test. The corresponding homogeneous groups were determined at significance level $P < 0.05$ and are indicated by a and b.

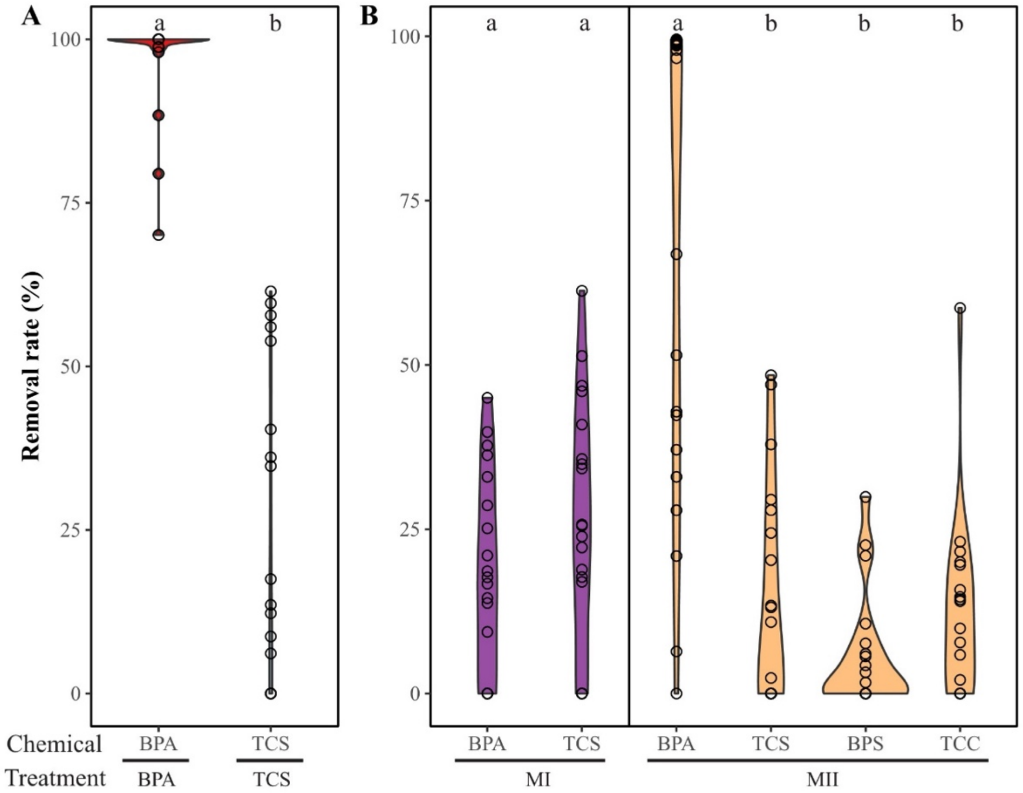

**Figure S6.** Non-metric multidimensional scaling (NMDS) based on Bray-Curtis dissimilarity matric illustrating the beta-diversity (between-sample differences in community composition) of the initial inoculum, the controls, and the treatments. The analysis was done using the exact sequence variants generated from DADA2 pipeline (Callahan et al., 2017). Raw sequences were processed using the DADA2 pipeline according to the DADA2 tutorial (v1.14) in R. The primers of sequences were trimmed using Cutadapt, and the resulting sequences were quality filtered with customized modifications as follows: truncLen=c(220, 190), maxEE=2, truncQ=2, maxN=0, rm.phix=TRUE, trimLeft=c(0,0). Subsequently, denoising, merging and chimera removal were completed according to the DADA2 pipeline tutorial. All sequences were aligned and assigned taxonomically using the SILVA v.132 reference database. The $M^2$ value and *P*-value were obtained from Procrustes statistic: $M^2$ value represents sum of squared deviations between sample pairs, and the lower value mean a better fit between matrices.

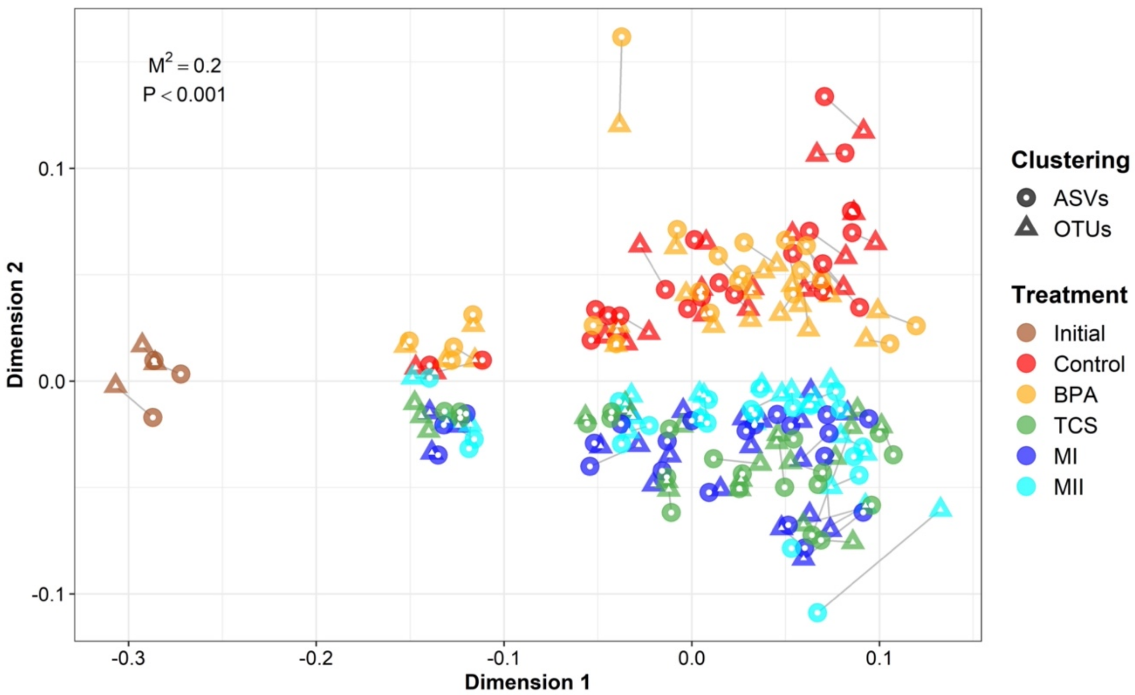

**Figure S7**. Bacterial community dissimilarity between the controls and micropollutant-treated microcosms (A-D) and the degree of variances between slopes indicated by |t| values (E). In A-D panels, the slopes (*v*) indicate rates of divergent succession of the micropollutant-treated communities from the control communities, estimated with a linear regression fitting between treatment-versus-control distance and time. In E panel, the significance of each linear regression is tested by permutation (1000 iterations), and significance level was determined at $P < 0.05$. The significant variances between slopes are estimated by bootstrapping (1000 iterations), followed by pairwise t-test with Bonferroni correction ($P < 0.05$).

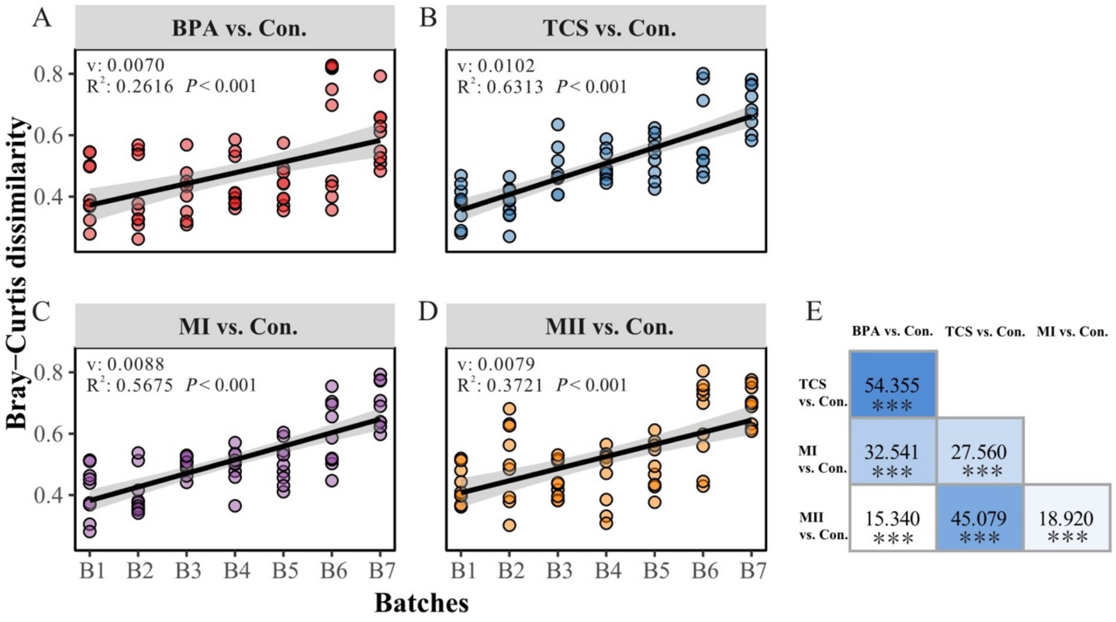

**Figure S8**. Relative abundance of the sensitive and tolerant taxa in each microcosm batch. Within each ecological group, the relative abundance was presented by the sum of the OTUs assigned to that group. Three growth progression phrases across all microcosm batches were identified based on similar average gene copy numbers and the time continuity of the inoculation phase 1 (B1–B2), phase 2 (B3–B4), and phase 3 (B5–B7).

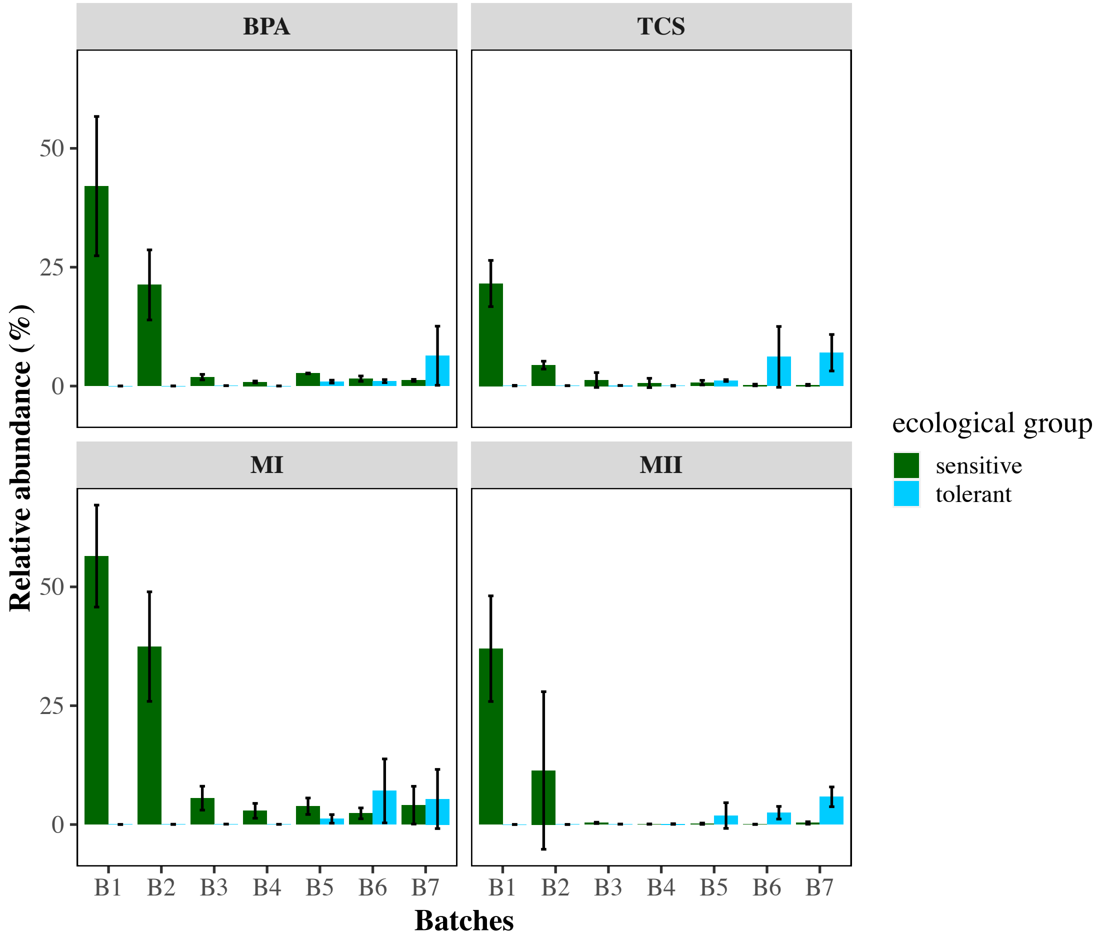

**Figure S9** Venn diagram illustrating the unique fraction of bacterial OTUs assigned to ecological categories found in the treatments but not shared in the controls when analyzed with the same filtering criteria as done for ecological grouping. The percentage in each sector of the Venn diagram presents the proportion of OTUs that are unique to either the controls or treatments, or that are shared between the two. A large fraction of the bacterial OTUs was unique to each of three ecological categories. This suggests that the identification of ecological strategies was not biased towards the selection of bacterial taxa whose high abundance in each defined phase was due to simple enrichment in the controls over time.

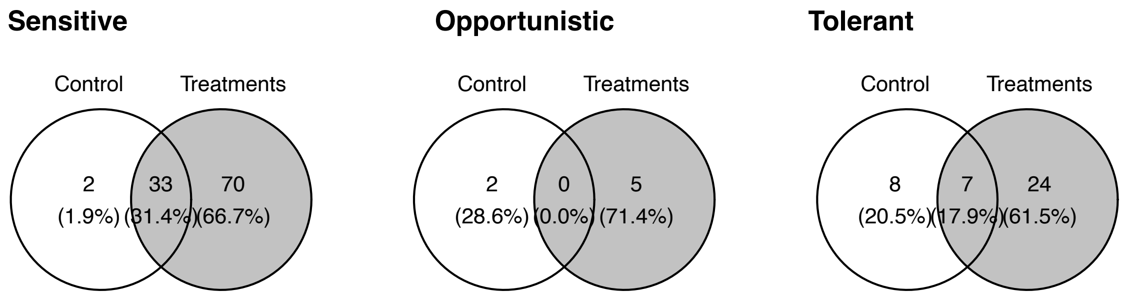
